# CAG repeat expansions create splicing acceptor sites and produce aberrant repeat-containing RNAs

**DOI:** 10.1101/2023.10.16.562581

**Authors:** Rachel Anderson, Michael Das, Yeonji Chang, Kelsey Farenhem, Ankur Jain

## Abstract

Expansions of CAG trinucleotide repeats cause several rare neurodegenerative diseases. The disease-causing repeats are translated in multiple reading frames, without an identifiable initiation codon. The molecular mechanism of this repeat-associated non-AUG (RAN) translation is not known. We find that expanded CAG repeats create new splice acceptor sites. Splicing of proximal donors to the repeats produces unexpected repeat-containing transcripts. Upon splicing, depending on the sequences surrounding the donor, CAG repeats may become embedded in AUG-initiated open reading frames. Canonical AUG-initiated translation of these aberrant RNAs accounts for proteins that are attributed to RAN translation. Disruption of the relevant splice donors or the in-frame AUG initiation codons is sufficient to abrogate RAN translation. Our findings provide a molecular explanation for the abnormal translation products observed in CAG trinucleotide repeat expansion disorders and add to the repertoire of mechanisms by which repeat expansion mutations disrupt cellular functions.

## Introduction

Expansions of CAG trinucleotide repeats are associated with at least twelve degenerative disorders, including Huntington’s disease (HD), dentatorubral-pallidoluysian atrophy (DRPLA), spinal and bulbar muscular atrophy (SBMA), and several spinocerebellar ataxias (SCAs)^1,2^. In each of these diseases, the associated gene is polymorphic for the number of CAG repeats, and disease manifests when the repeat number exceeds a certain threshold^1^. In most CAG repeat expansion disorders (including HD, and SCA types 1, 2, 3, 6, 7, and 17, DRPLA, and SBMA), the CAG repeat tract is located in the protein coding region, and codes for a polyglutamine stretch^1^. An expanded polyglutamine tract renders proteins aggregation prone, and this repeat-dependent protein aggregation contributes to disease pathology^1^.

Besides encoding for polyglutamine-containing proteins, CAG repeat expansions can produce cellular dysfunction via at least two additional routes, even when they occur outside of the canonical protein coding regions. First, repeat-containing RNAs can agglomerate in the nucleus as pathogenic foci^3,4^. RNA foci result from multivalent intermolecular base-pairing interactions templated by the GC-rich repeat tract^5^. These foci sequester various RNA binding proteins, and cause widespread RNA processing defects^6^. Second, repeat-containing RNAs undergo translation without requiring an identifiable AUG start codon, in a process known as repeat-associated non-AUG (RAN) translation^7–10^. RAN translation produces repeat-containing proteins in multiple reading frames, which may form protein aggregates, and are potentially lethal to the cell^11^. The molecular mechanism of RAN translation is not entirely clear.

In our earlier work, we showed that RNAs that consist primarily of expanded CAG trinucleotide repeats, which are not present in a canonical open reading frame (ORF), accumulate at nuclear foci. These repeat-containing RNAs did not undergo RAN translation even when the repeat number was increased to >400 tandem CAGs^5,12^. RAN translation is reported to be influenced by the sequences adjacent to the repeat tract^10,13^. To assess the role of flanking sequences, we previously generated a library of CAG repeat-containing plasmids where various 250 nucleotide long sequences were cloned upstream of 240×CAG repeats^12^. We found that a subset of these flanking sequences led to RAN translation of the CAG repeat, while others were still retained at nuclear foci^12^. We were not able to identify sequence motifs in the upstream flanking sequences that differentiated the two classes.

Here, we set out to identify the features of the flanking sequences that determine whether a given CAG repeat-containing RNA will be sequestered at nuclear foci or would undergo RAN translation. We hypothesized that an expanded CAG repeat itself could provide new splicing acceptor site(s), and produce aberrant repeat-containing transcripts. Our hypothesis was guided by three observations. One, livecell RNA imaging experiments showed that RNAs that undergo RAN translation are first retained at nuclear foci for a prolonged period where they colocalize with the splicing machinery^12^. Two, mammalian 3’ splice sites have a nearly invariant ‘AG’ dinucleotide that marks the splice junction (minimal splice acceptor represented as Y_6-12_NAG**NNN**, where Y_6-12_ indicates the upstream pyrimidine rich tract, N is any nucleotide, and the bold bases mark the exon)^14^. In fact, at least 141,000 (∼64%) of annotated human splicing acceptor sites harbor a CAG trinucleotide at splice junctions, and >1400 of these sites exhibit two or more (up to nineteen) consecutive CAG trinucleotides (Supp. Fig. 1A). Three, CAG repeat expansions induce transcriptome-wide splicing changes^6,15^, and in some cases, are reported to induce missplicing of the gene harboring the expanded repeat^16–18^.

**Figure 1.**
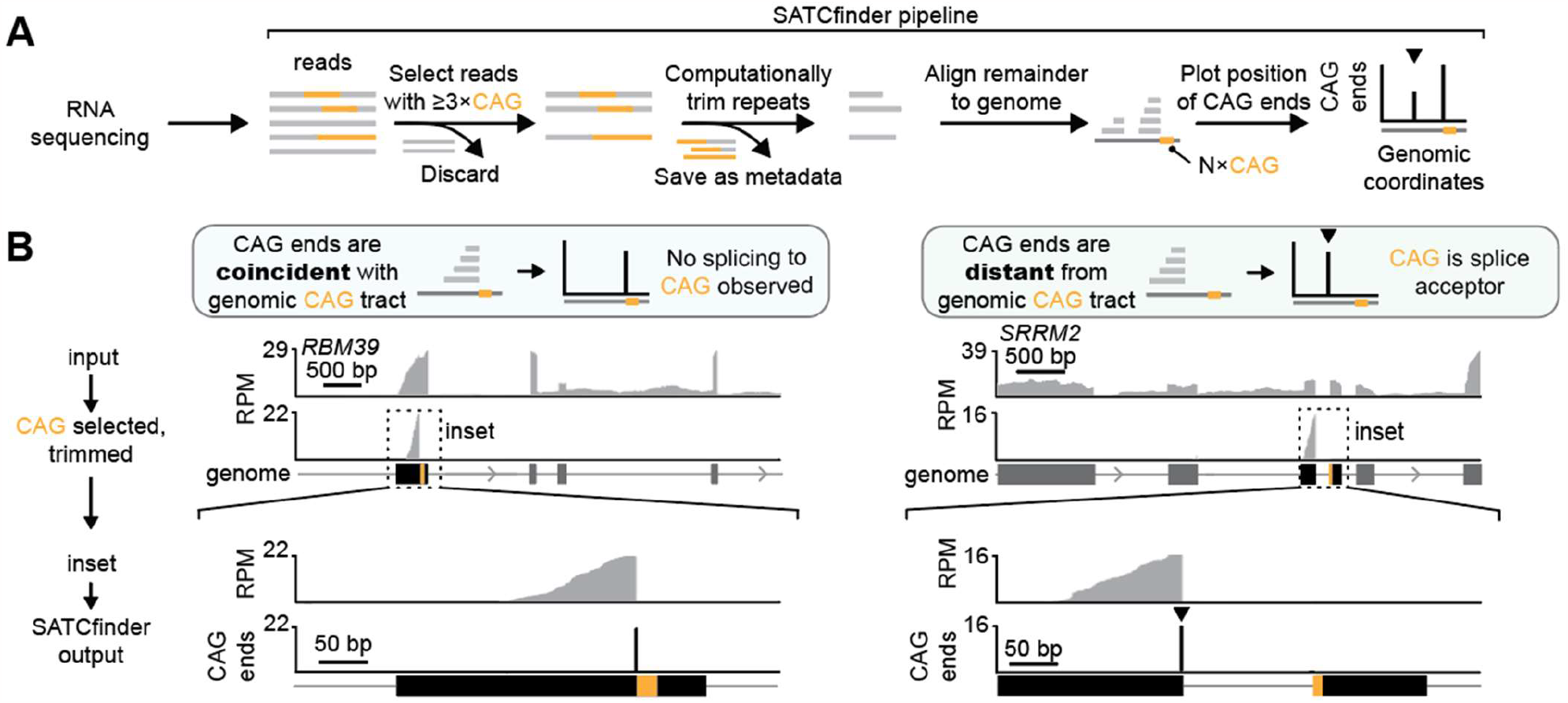
A. Schematic for SATCfinder. Reads with ≥3xCAG/CTG repeats are selected. The CAG repeats are computationally removed and these trimmed reads are aligned to the genome. The genomic coordinates of the base immediately before the repeat (CAG end) in the trimmed and mapped reads is tracked. SATCfinder outputs the number of CAG ends per million mapped reads at a given genomic coordinate. The peak at the repeat reflects reads where CAG repeats are not a part of the splice junction, while a distal upstream peak (at the nearest upstream exon, typically within a few kb) indicates the location of the splicing donor (marked by an arrowhead ▾). **B**. Representative genes comparing standard RNA sequencing analysis to SATCfinder output.

GC-rich repeat-containing sequences are difficult to analyze using standard RNA sequencing approaches, and thus, aberrantly spliced repeat-containing transcripts may have been missed in previous analyses. Thus, we developed an analysis pipeline that specifically captures splicing events at tandem CAG repeats. We find that cognate and near-cognate splicing donors in the surrounding sequences can splice into an expanded CAG repeat tract. This splicing is dependent on the number of CAG repeats, and can place the repeats in unexpected sequence contexts. Interestingly, disease-causing CAG repeat expansions, which are proposed to undergo RAN translation, produce RNAs that harbor CAG repeats in AUG-initiated ORFs. Disruption of the relevant splicing donor site or the AUG initiation codon is sufficient to abrogate the production of proteins that are attributed to RAN translation. Our findings provide a cogent mechanistic explanation for how expanded CAG repeats produce abnormal protein products and suggest a novel pathomechanism by which repeat expansions can interfere with RNA processing and cellular functions.

## Results

### Sequence analysis pipeline for detecting splicing to CAG repeats

Analysis of repeat-containing RNAs using short-read sequencing technologies is challenging. GC-rich repeats are difficult to reverse transcribe and amplify, and these sequences are substantially under-represented in standard RNA sequencing libraries^19,20^. The commonly used sequence alignment pipelines also exclude these regions, as repeat-containing reads may map to multiple locations in the genome, or because the reference genome lacks an expanded repeat tract^21^. To address the first issue, we optimized the library preparation protocol, notably by using a group II intron reverse transcriptase that can reverse transcribe structured^22^ and GC-rich^23,24^ RNAs with high fidelity (see Methods). To address the issue of aligning repetitive reads, we developed a pipeline, SATCfinder, to identify and visualize splicing at tandem CAG repeats in RNA sequencing data (Fig. 1A). In brief, SATCfinder’s input is paired end RNA sequencing data. All reads that contain ≥3×CAG/CTG repeats (to capture both sense and antisense transcripts) and at least 15 additional bases that are not a part of the CAG repeat (to allow appropriate alignment and junction identification) are selected. The CAG repeats are computationally trimmed and the remaining non-repetitive portion of the reads, together with the paired read mates, are aligned to the reference genome (see Methods). The mapping coordinate of the CAG end of the trimmed reads (hereafter, CAG end) is tracked. The number of CAG ends coinciding with the genomic coordinate of the CAG repeat versus those aligning farther away to upstream splicing donors allows us to quantitatively assess the extent of splicing to the repeat tract (Fig. 1A). For example, when CAG repeats are not a part of the splicing acceptor (i.e., they are contained entirely in an exon or an intron), the position of the CAG repeats in mRNA matches the location of the repeat tract in the genome. For a representative gene in this class, *RBM39*, the CAG ends in RNA identified using SATCfinder coincide with the end of the CAG repeats in the genome (in 99.5% of reads, Fig. 1B left, Supp. Fig. 1B). Likewise, we examined several other genes where CAG repeats are annotated to be in the middle of an exon, and in each case, >99% of CAG ends mapped immediately adjacent to the CAG repeat tract in the genome (Supp. Fig. 1B, D).

Sixty-eight human genes are annotated to have ≥3×CAG at splice acceptor sites (Supp. Fig. 1A, Supplemental Table 1). For this class, the non-repetitive sections of CAG-containing reads map to a location upstream of the CAG repeat in the genome (see representative example *SRRM2*, Fig. 1B, right). A vast majority (∼95%) of CAG ends align to the annotated splicing donor site upstream from the tandem CAG acceptor site (Fig. 1B, right, Supp. Fig. 1C, E). The remainder (∼5%) of CAG ends coincide with the repeat tract in the genome, likely arising from unspliced pre-mRNAs (Fig. 1B, right, Supp. Fig. 1C). Treatment with a splicing inhibitor increased the proportion of reads corresponding to unspliced CAG repeat-containing RNAs by ∼10-fold (from 3% to 33% of CAG ends on treatment with 25 nM pladienolide B, Supp. Fig. 1F). Similar results are observed for other genes in this class (Supp. Fig. 1C-D). Altogether, these results demonstrate that our library preparation protocol coupled with SATCfinder allow us to examine events where CAG repeats form splicing acceptor sites.

### CAG repeat expansions result in mis-splicing of repeatcontaining RNAs

We then used SATCfinder to examine whether CAG repeat expansions could induce mis-splicing of repeat-containing RNAs. We previously generated a small library of sequences where 240×CAG repeats were cloned downstream of various 250 nucleotide flanking sequences^12^ (Fig. 2A). In these constructs, multiple stop codons were incorporated immediately upstream of the CAG repeats to eliminate translation readthrough from any upstream ORFs into the repeats. In about half of these constructs, the CAG repeat-containing RNA was retained in the nucleus at foci, while in other cases, the repeat-containing RNA was exported to the cytoplasm, underwent RAN translation, and formed perinuclear aggregates (Fig. 2A). Cytoplasmic localization also coincided with substantial cellular toxicity^12^.

**Figure 2.**
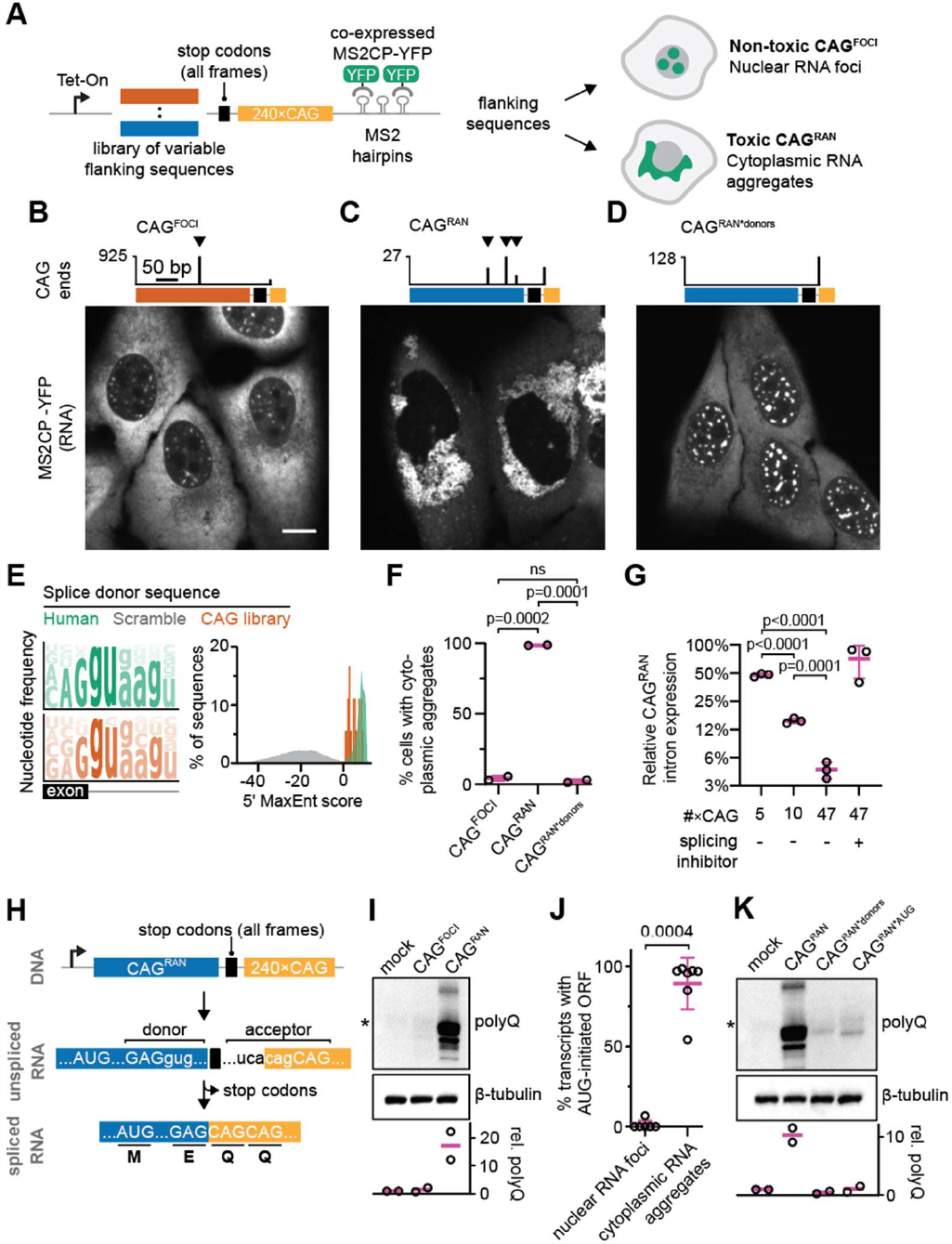
A. Schematic for constructs with 240×CAG repeats with a variable 250-base flanking sequence^12^. Some flanking sequences result in retention of the repeat-containing RNA in the nucleus while others induce RAN translation and cell toxicity. **B, C, D**. *Top*, SATCfinder output for representative CAG constructs, where the x-axis is the base coordinate within the flanking sequence region and the y-axis indicates the number of CAG ends per million mapped reads. *Bottom*, Representative fluorescent images of cells expressing the indicated constructs. Micrographs are representative of ≥ 2 independent experiments. Scale bar depicts 10 μm. **E**. *Left*, sequence logos for 5’ splice sites annotated in the human genome and those observed in the CAG flanking sequence library, where the x-axis indicates the position within the 9-base donor sequence and the letter height depicts the probability of observing the base. *Right*, 5’ MaxEnt scores for 220419 annotated human splice donors, all 262144 possible randomly generated 9-mers, and the 20 detected splice donors in the CAG flanking sequence library. **F**. Quantification of the percentage of cells with cytoplasmic RNA aggregates by fluorescence microscopy. Each data point represents an independent experiment with ≥ 500 cells per experiment, and are summarized as mean ± SD. **G**. Real-time quantitative PCR quantification of the relative expression of CAG^RAN^ intron normalized to the expression of the 5’ end of the CAG^RAN^ transcript. Data show the mean ± SD for three independent RNA isolations. **H**. Schematic for CAG^RAN^, depicting the transcription initiation site (as a right-facing arrow), flanking sequence, and 240×CAG repeats intervened by stop codons in each frame. The sequence of a representative donor that splices to the CAG tract is shown, with bases in the exon in uppercase. After splicing, the stop codons are removed and the CAG repeat is embedded in an AUG-initiated ORF. **I**. Immunoblot for cells expressing CAG^FOCI^ and CAG^RAN^ using a polyglutamine antibody. An endogenous protein (TBP), marked by an asterisk, is also detected by this polyglutamine antibody^10,57^. **J**. Percentage of repeatcontaining transcripts where the repeats are observed in AUG-initiated ORFs for constructs that produce RNA foci only or exhibit RAN translation. Each data point is one construct. **K**. Immunoblot for the indicated samples using a polyglutamine antibody. CAG^RAN*donors^ has point mutations at all identified splice donor sites; CAG^RAN*AUG^ has point mutations at two AUGs. Immunoblots are normalized first to tubulin, then to the parent cell line without a repeat-containing construct (mock). Immunoblots and quantification of relative polyglutamine abundance (as mean ± SD) are representative of ≥ 2 independent experiments. Significance values in F, G, and J are calculated using Student’s t-test. MS2CP-YFP: bacteriophage MS2 coat protein tagged with YFP.

We performed RNA sequencing on cells expressing the various 240×CAG repeat-containing RNAs that either form nuclear foci (6 cell lines, with CAG^FOCI^ as a representative of this class, full sequences are provided in Supplemental Table 2) or undergo RAN translation (7 cell lines, with CAG^RAN^ as a representative of this class). Our analysis using SATCfinder revealed that the CAG repeats in RNA were frequently stitched to regions in the upstream flanking sequences, and the intervening region had been removed (see representative examples for CAG^FOCI^ and CAG^RAN^ in Fig. 2B-C, top; and other examples in Supp. Fig. 2A-B).

Several lines of evidence indicate that the junctions identified by SATCfinder arise from splicing to CAG repeats. One, quantitative and end-point PCR on the genomic DNA and RNA confirmed that the desired sequences were integrated into the genome but the intervening sequences were only removed from the RNA (Supp. Fig. 3A-C). Two, the read junctions identified by SATCfinder harbor signatures of splicing donors (Fig. 2E, left, Supp. Fig. 3D). We evaluated these identified donor sites using a computational splicesite prediction algorithm, MaxEntScan^25^. This algorithm reports a log-odds ratio of observing a motif in true versus decoy splice donor sites, where a higher 5’ MaxEnt score indicates a stronger putative donor site^26^. Known human splice donors score in the range 0 to 12 (>95% score above 2; median 8.6, Fig. 2E, right). The donors that were identified by SATCfinder to splice to CAG repeats in our library had positive 5’ MaxEnt scores (1.59 – 9.40, median 4.95; Fig. 1E, Supp. Fig. 3D), indicating that these sequences are canonical splice donors. Three, chemical inhibition of splicing using pladienolide B^27^ led to a dose-dependent reduction in the frequency of splicing from the upstream flanking region to the CAG repeats (Supp. Fig. 3C, E). Finally, point mutations to the crucial **G**GU motifs of the donors (**G**GU→**G**GA) were sufficient to eliminate splicing to the repeats (Fig. 2D, Supp. Fig. 3F).

**Figure 3.**
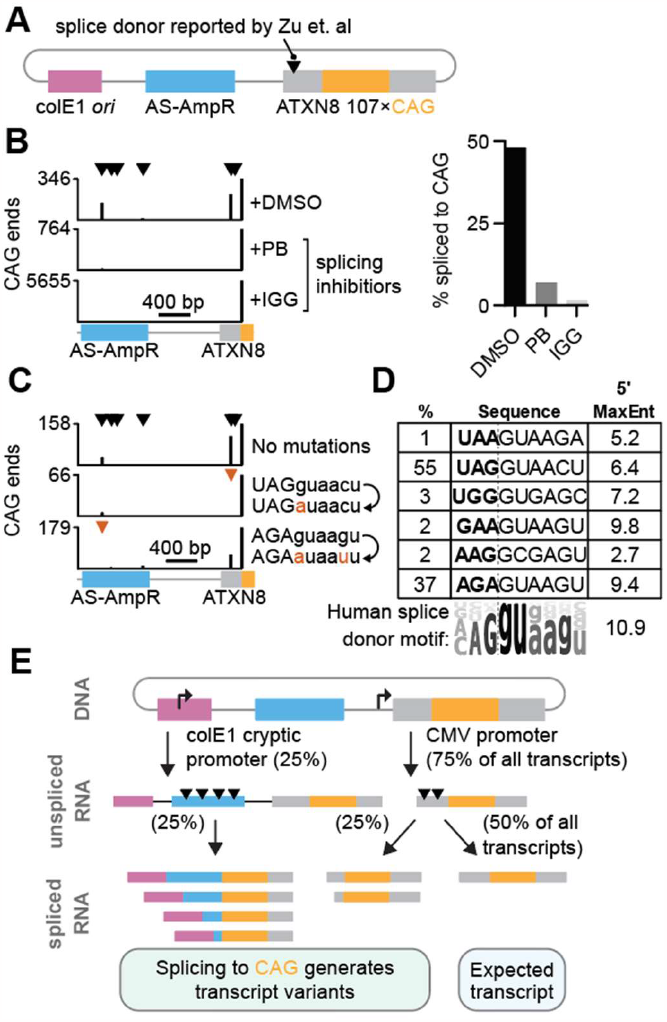
A. Schematic for the ATXN8 mini-gene expressing ∼100 bp of endogenous *ATXN8* sequence directly upstream from 107×CAG repeats^10^. The ampicillin resistance gene (AS-AmpR) and colE1 origin of replication are indicated. **B**. *Left*, SATCfinder output for cells transfected with the ATXN8 construct in the presence of splicing inhibitors 25 nM pladienolide B (PB), or 15 μM isoginkgetin (IGG), or 0.1% DMSO (DMSO) as control. *Right*, quantification of the % of CAG ends that reflect splicing to the repeat. **C**. Similar to B but for ATXN8 constructs without or with point mutations to identified donor sites. **D**. Sequences of the identified splicing donors in the ATXN8 construct with corresponding percentage of reads arising from each donor. The sequence logo for the consensus human splice donor is presented for comparison. **E**. Schematic for the various CAG repeat-containing transcripts produced upon splicing from the ATXN8 construct.

Splicing was dependent on the number of CAG repeats, and its frequency progressively increased with the repeat number (Fig. 2G). Similar splicing patterns were observed across various cell lines (Supp. Fig. 3G). A variety of donor sequences could be spliced to the CAG repeats, and the consensus sequence of the donors identified in our flanking sequence library closely resembles the consensus sequence for the annotated donor sites in the human genome (Fig. 2E, Supp. Fig. 3H-I). In summary, these results show that splicing from putative donors, occurring by chance in our library of flanking sequences, resulted in unexpected CAG repeat-containing RNAs that differ from the intended sequences that were integrated in the genome.

While we engineered stop codons in each frame immediately upstream of the CAG repeats, a subset of these constructs produced polyglutamine-containing proteins (Fig. 2H-I). Splicing to CAG repeats could create mature transcripts where the proximal stop codons are spliced out and the repeats are placed within AUG-initiated ORFs. To test this idea, we assembled the full-length mature mRNAs produced from these constructs using our RNA sequencing data. In all cases where we observed translation of the repeat region, we found AUG-initiated ORFs, generated after splicing, that contain the repeat region (Fig. 2J, Supp. Fig. 4A). On the other hand, sequences that formed nuclear foci either did not exhibit substantial splicing to CAG repeats or, upon splicing, did not produce AUG-initiated ORFs that contain the CAG repeats (Fig. 2J, Supp. Fig. 4A). Across the various cell lines, the abundance of RNAs with repeat-containing AUG-initiated ORFs correlated with the levels of polyglutamine protein produced (Pearson’s correlation coefficient, r = 0.83, between polyglutamine immunofluorescence and RPM of transcripts carrying an AUG-initiated ORF, Supp. Fig. 4B-C).

**Figure 4.**
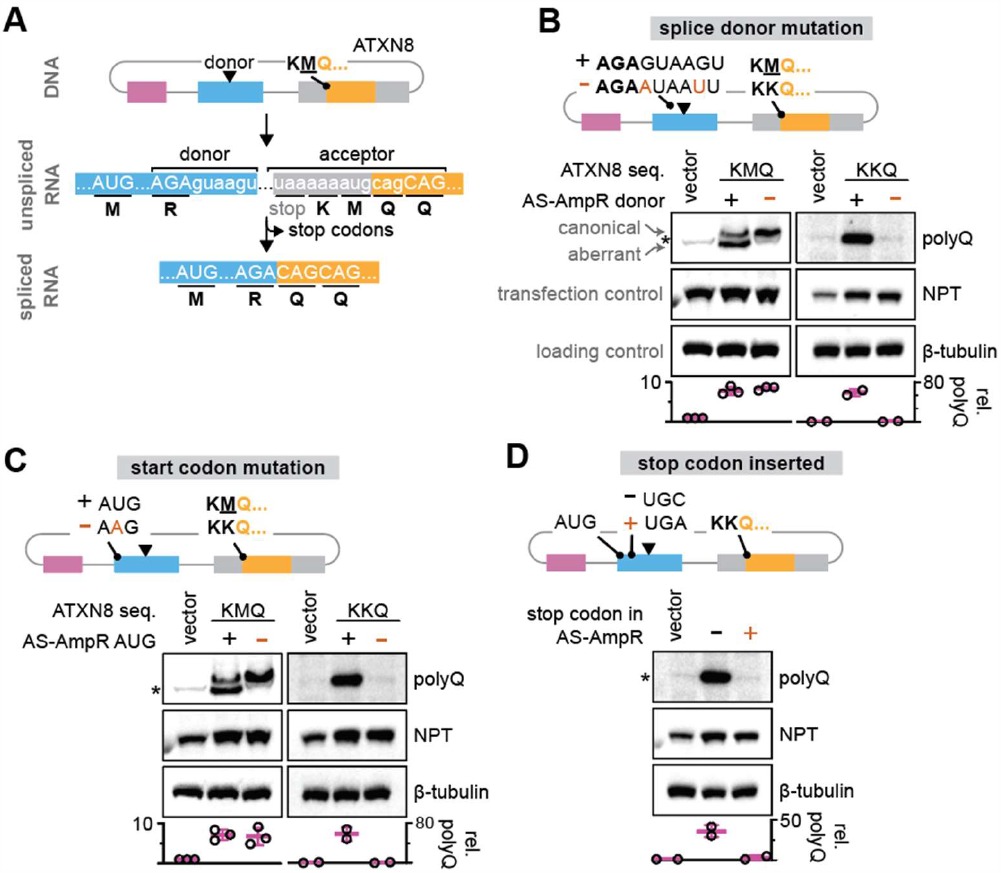
A. Schematic for ATXN8 mini-gene. Upon splicing, the upstream stop codon is removed and the CAG repeat is embedded in an AUG-initiated ORF. **B, C, D**. Immunoblots from cells expressing the indicated ATXN8- and ATXN8^KKQ^-derived constructs that interrupt the predicted ORF by mutating the splice donor (B), mutating the identified in-frame AUG initiation codon (C), or by introducing a stop codon (D). Band intensities are normalized first to NPT (neomycin phosphotransferase), expressed in *cis* from the plasmid, and then to the endogenous protein (TBP, marked with an asterisk) in the control transfected with a similar vector but encoding for GFP (vector). Tubulin is included to show equivalent loading between conditions, but is not used for normalization due to potential variations in transfection efficiency. Immunoblots and quantification of relative polyglutamine abundance (as mean ± SD) are representative of ≥ 2 independent transfections.

We chose one representative sequence, CAG^RAN^, to further examine the role of splicing in the translation of the repeat region. Mutation of the identified splice donor sites in CAG^RAN^, which eliminated splicing to the repeat tract, also abrogated the production of aberrant polyglutamine proteins (Fig. 2K). The mature repeat-containing transcripts produced from CAG^RAN^ contained an ORF with two in-frame AUG codons (Fig. 2H). Disruption of these AUG codons by single-base mutations at each site similarly eliminated the polyglutamine product (Fig. 2K). N-terminal sequencing of the polyglutamine protein produced in CAG^RAN^ confirmed that it is methionine-initiated, and the first six amino acids in this protein (MRFLAT) are as expected for translation initiation from the first AUG in the ORF (Supp. Fig. 4D). Interestingly, pharmacological inhibition of splicing led to the retention of the repeat containing RNA in the nucleus at foci (Supp. Fig. 5A-B). Likewise, point mutations to the splice donors in CAG^RAN^ were sufficient to convert this sequence to the foci-forming class (Fig. 2D, bottom, and Fig. 2F). Unlike CAG^RAN^, expression of this sequence with mutated donors (CAG^RAN*donors^) did not induce cell toxicity (Supp. Fig. 5C), consistent with our earlier report that expression of repeat-containing RNAs that only produce nuclear foci, do not induce overt cell death^12^. Thus, the aberrant translation products in CAG^RAN^ result from canonical AUG-initiated translation of aberrant transcripts that arise from splicing to expanded CAG repeats, and disruption of splicing or the appropriate AUG start codons is sufficient to eliminate the translation of the repeat tract.

**Figure 5.**
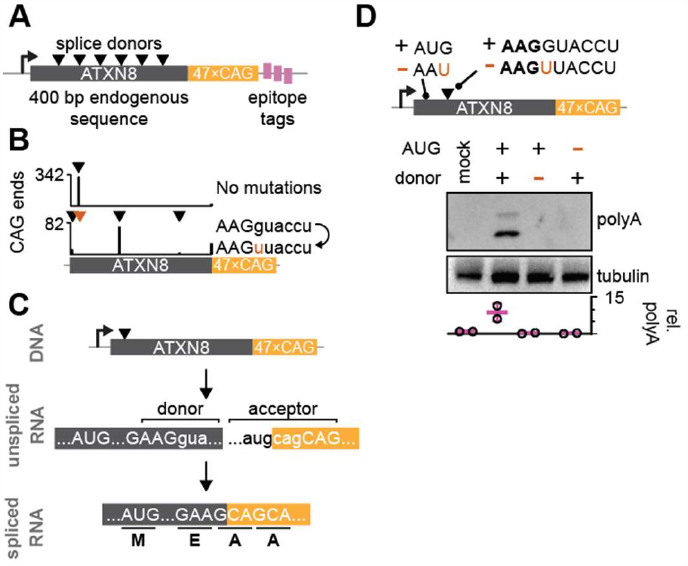
A. Schematic for design of ATXN8 mini-gene with 400 bases of endogenous ATXN8 sequence fused directly to 47×CAG repeats, followed by epitope tags in each reading frame. **B**. SATCfinder output for ATXN8 constructs with native upstream sequence, without or with a point mutation to the predicted donor site. **C**. Schematic for ORF resulting from ATXN8 mini-gene. Upon splicing, the upstream stop codon is removed and the CAG repeat is embedded in an AUG-initiated ORF. **D**. Immunoblot for the indicated samples using an anti-HA antibody. The HA epitope is in the polyalanine frame. Immunoblots are normalized first to tubulin, then to the parent cell line without a repeat-containing construct (mock). Immunoblots and quantification of relative HA abundance (as mean ± SD) are representative of ≥ 2 independent experiments.

### CAG repeats with native disease-associated flanking sequences form splice acceptors

Besides an acceptor site, splicing requires cis-acting elements, such as a polypyrimidine tract, that facilitate spliceosome assembly. In our flanking sequences library, CAG repeats were placed in a synthetic sequence context which likely provided the polypyrimidine tract (sequence in Supplemental Table 2). Thus, we next investigated whether disease-associated CAG repeats in their native upstream sequence context may also act as splicing acceptors. We focused on disease models where expanded CAG repeats are reported to produce aberrant RAN translation products.

The first report on RAN translation of CAG repeats utilized a minigene with 107×CAG repeats flanked by sequences native to the *ATXN8* gene^10^, associated with SCA8 (Fig. 3A). Interestingly, Zu et al. also observed splicing from an upstream splice donor to the CAG repeats, but the resulting spliced transcript did not contain ORFs that explained the observed aberrant translation products (Fig. 3A). To investigate whether there were additional donor sites, we transfected HEK293T cells with this ATXN8 minigene, similar to the previous study, and analyzed the resulting RNA using SATCfinder. Our analysis revealed that only about half of the CAG-containing reads were unspliced and mapped to the expected mini-gene sequence (Fig. 3B). The remaining CAG-containing reads mapped to distant sites on the plasmid, suggesting that additional mature CAG-repeat-containing RNAs are produced in these cells (Fig. 3B). Using a conservative threshold of ≥100 unique reads supporting a putative splice donor, we identified six donor sites (Fig. 3B, Supp. Fig. 6A). The donors contain the nearly invariant ‘**G**GU’ motif found in human splice donors, and have positive 5’ MaxEnt scores (2.7 – 9.82, Fig. 3D). Splicing inhibition using pladienolide B or isoginkgetin^28^ significantly decreased the number of reads that reflect splicing to the repeat tract (84% and 98% reduction in splicing for 25 nM pladienolide B and 15 μM isoginkgetin treatments, respectively, Fig. 3B). Point mutations to the donor sites completely abolished splicing from the respective donors (Fig. 3C), further validating that these donors are spliced to the CAG repeat tract. Similar results were observed for other CAG-containing mini-genes where the proximal sequences are derived from the genes *HTT, JPH3, ATXN3*, and *DMPK-AS*, associated with the diseases HD, Huntington disease-like 2, SCA3, and myotonic dystrophy type 1 respectively (Supp. Fig. 6B).

**Figure 6.**
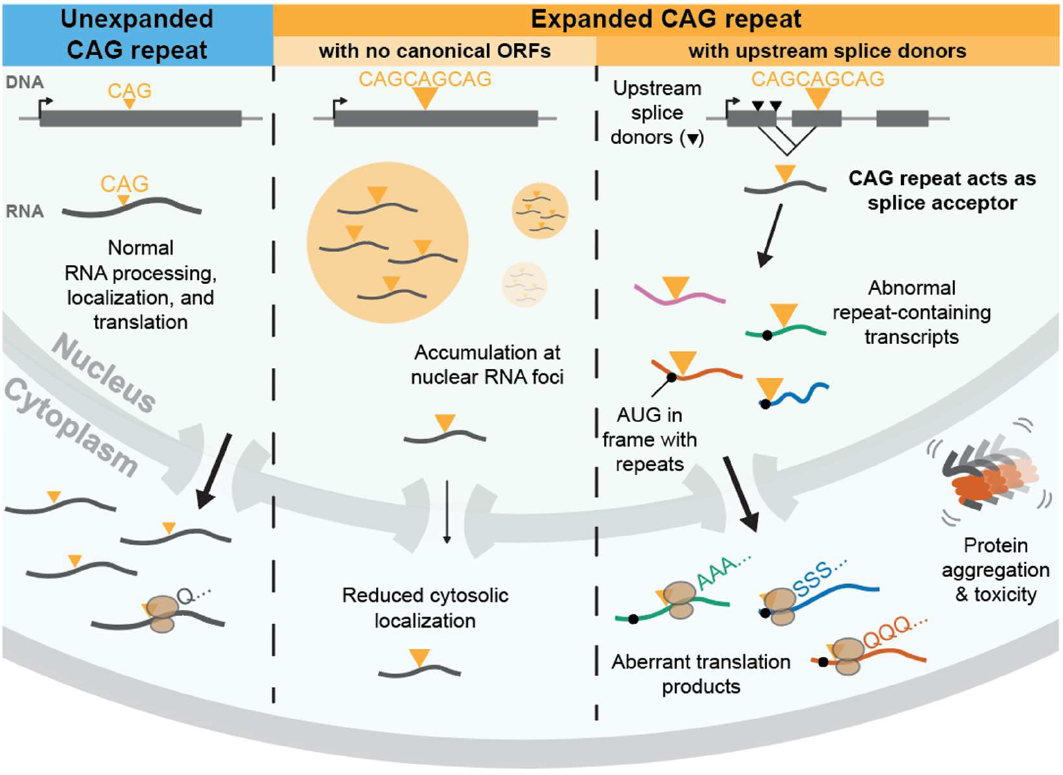
Model for the sub-cellular localization of RNA with expanded CAG repeats. In the absence of an expanded repeat, the RNA is normally processed and exported to the cytoplasm. If the RNAs contain expanded CAG repeats outside of an ORF, the RNAs are retained at nuclear foci, where they sequester splicing factors. If the CAG repeats are located downstream of potential splice donors, the donors may be spliced to the CAG repeat in a repeat number dependent manner. Splicing generates new RNA isoforms where the CAG repeat may be present in AUG-initiated ORFs. Translation of these AUG-initiated repeat-containing ORFs produces aberrant homopolymeric proteins that may aggregate and contribute to cellular toxicity.

We examined the origin of the splicing donor sequences in these mini-genes, and to our surprise, we found that a significant fraction of the spliced reads arose from a donor located in the ampicillin resistance cassette more than 1 kilobase upstream from the CAG repeats (Fig. 3B). The AmpR gene is encoded in the antisense direction (AS-AmpR) with respect to the CAG repeats. The bacterial AmpR promoter should not initiate transcription in mammalian cells nor would it produce CAG repeat-containing transcripts (see full sequence map in Supp. Fig. 6C). To determine how a donor in the AS-AmpR sequence could splice to the CAG repeats, we examined the raw (non-CAG-selected) read alignments. We noted a region of high coverage beginning within the colE1 *E. coli* origin of replication and continuing through the AS-AmpR region (Supp. Fig. 6D). This coverage did not result from readthrough of the polyadenylation signal from the adjoining neomycin resistance gene (Supp. Fig. 6D). Transcript assembly using StringTie^29^ confirmed that the AS-AmpR transcript originated within the colE1 region (Supp. Fig. 6E). Reads corresponding to this region were also observed in other plasmids that harbor colE1 origin of replication (Supp. Fig. 6F), and the transcripts initiating from colE1 were ∼30% as abundant as those from the SV40 and CMV promoters (Supp. Fig. 6G). This observation is consistent with reports that the colE1 origin of replication acts as a cryptic promoter in eukaryotic cells and may result in spurious unintended transcripts in cells transfected with plasmids carrying this bacterial sequence^30,31^. Thus, transcription from the cryptic colE1 promoter and splicing from donor sites in the AS-AmpR region to the repeat tract gives rise to mature transcripts containing CAG repeats embedded in various 5’ sequence contexts (Fig. 3E). Taken together, these results demonstrate that an expanded CAG repeat tract flanked by sequences native to the disease-causing gene can act as splice acceptor sites, and splicing from upstream donors may result in unexpected repeat-containing transcripts.

### Canonical translation of aberrantly spliced CAG-repeat-containing RNAs accounts for RAN products

We then investigated whether the spliced RNAs produced from disease associated mini-genes with repeat-containing ORFs might explain the production of spurious repeat-containing proteins. We examined the ATXN8 mini-gene, where the CAG repeat is immediately preceded by an AUG codon in the polyglutamine frame (Fig. 4A). Transfection of the ATXN8 plasmid in HEK293T cells resulted in the production of multiple polyglutamine-containing proteins (Fig. 4B), as has been previously reported^10^. Prior work demonstrated that mutation of the in-frame start codon immediately preceding the CAG repeats (construct ATXN8^KKQ^, Fig. 4A) eliminated one of the polyglutamine products, while other polyglutamine-containing proteins produced from this plasmid are not affected. These products have been attributed to non-canonical RAN translation of the repeat region^10^.

We found that these plasmids produced several spliced RNAs with AUG-initiated ORFs that contain the CAG repeats (Fig. 4A; see other ORFs in Supp. Fig. 7A). One of the major transcripts with an ORF in the polyglutamine frame arises due to splicing from a donor in the AS-AmpR cassette. Point mutations to this donor in AS-AmpR, ∼1800 bases upstream from the CAG repeats, that abolished splicing at this site (Fig. 2C), also eliminated the aberrant polyglutamine product in both ATXN8 as well as ATXN8^KKQ^ constructs (Fig. 4B). Splicing from this donor to CAG repeats creates an ORF with an AUG start codon located 135 nucleotides upstream from the donor site. Again, a single mutation to this AUG site, located in AS-AmpR, completely abolished the aberrant polyglutamine protein without affecting the canonical translation product (Fig. 4C). As an additional test for our model, we interrupted the AUG-initiated ORF with a single stop codon. This single-base mutation in the AS-AmpR region to generate a stop codon likewise eliminated the aberrant polyglutamine product (Fig. 4D). Finally, we deleted a single nucleotide 10 bases upstream from the donor site, which would shift the translation frame for CAG repeats from polyglutamine to polyserine. Consistent with this expectation, the single-base deletion abrogated aberrant polyglutamine production with a concomitant ∼500% increase in high-molecular weight polyserine-containing protein (Supp. Fig. 7B). Taken together, these results demonstrate that at least a subset of abnormal translation products arising from these mini-genes can be accounted for by splicing of upstream donors to CAG repeats followed by canonical AUG-initiated translation of the repeat-containing RNA. Even though the splicing donor sequences in these mini-genes used to model disease are non-native, our results suggest that disease-associated expanded CAG repeats, flanked by their native sequences, can act as splicing acceptors and potentially generate aberrant transcripts with repeat-containing ORFs.

### Splicing from an endogenous donor in *ATXN8* generates AUG-initiated ORFs

Our observations raise the possibility that a similar mechanism may account for spurious, out of frame protein products observed in disease, where the region upstream of the repeat tract may provide a splice donor. We examined the sequence upstream of CAG repeats in the *ATXN8* gene and found several potential splicing donor sites (5’ MaxEnt scores ≥ 1, Supp. Fig. 8A). We cloned 400 nucleotides of this native 5’ flanking sequence and placed it immediately upstream of 47×CAG repeats. Downstream of the repeats, we incorporated distinct epitope tags in each reading frame (Fig. 5A, sequence in Supplemental Table 2). Upon transducing this construct in U-2OS cells, we observed one major splice donor that underwent splicing to the CAG repeat (5’ MaxEnt = 8.49). More than 90% of CAG ends reflected splicing to the repeats from this specific site (Fig. 5B). Disruption of this donor by a single-base mutation eliminated splicing from this location (Fig. 5B). Interestingly, when we mutated the original donor site, several new donor sites in the flanking sequence were spliced to the CAG repeats (5’ MaxEnt score for these sites ranging from 1.76 to 5.17 Fig. 5B, Supp. Fig. 8A). This observation is similar to other reports indicating that mutation of a splice donor can result in the activation of cryptic donor sites^32^, and suggests that the CAG repeat sequence in *ATXN8* creates a strong splicing acceptor.

Splicing from the identified donor site in *ATXN8* creates an AUG-initiated ORF in the polyalanine frame (Fig. 5C). Polyalanine products have been observed in brain tissue from SCA8 patients^10^. In our synthetic construct, this spliced ORF encodes for an ∼7 kD polyalanine containing protein which is detected via an HA-tag encoded downstream of the repeats in the polyalanine frame (Fig. 5D). Point mutations to the splicing donor or to the relevant AUG initiation codon abrogate polyalanine protein production (Fig. 5D, Supp. Fig. 8B-C). Although polyserine products have also been reported in SCA8 patient brains^7^, we do not observe translation of the repeat in the polyserine frame in this model (Supp. Fig. 8B). The small fragment of the *ATXN8* gene that we cloned does not produce RNA with AUG-initiated ORFs in the polyserine frame. Nonetheless, further characterization of the 5’ end of the full-length *ATXN8* transcript may reveal splice donors that could account for additional aberrant translation products that are observed in disease.

## Discussion

The diseases caused by simple repeat expansions have diverse pathomechanisms that encompass dysfunction at DNA, RNA, and protein levels. Expanded CAG repeats in DNA form secondary structures that induce repeat instability during DNA replication and repair^33–36^. The repeat-containing RNAs may form inter-molecular base pairs and agglomerate in the nucleus to form foci that sequester essential RNA binding proteins^3,4^. When translated, the repeats produce aggregation-prone homopolymeric proteins^37–39^. Our work uncovers yet another route by which the disease-causing CAG repeats may trick the cellular machinery (see model in Fig. 6). Expanded CAG repeats may create new splicing acceptor sites. Nearby donors may splice into the repeats, and thus generate a variety of repeat-containing transcripts. These transcripts may contain AUG-initiated ORFs that encompass the repeat tract and produce unexpected, but canonically translated, polypeptides. This mechanism may account for a subset of abnormal polypeptides that are observed in various CAG repeat expansion disorders^40–42^.

Mechanistically, how the expanded CAG repeat tract induces mis-splicing of the repeat-containing transcript remains to be investigated. One possibility is that the expanded CAG repeats may provide a battery of alternative acceptor sites. More than 800 human genes contain tandem CAG repeats (≥2xCAG) at the 3’ splice sites^43^. In several cases, these repeats create alternative splice junctions, with the upstream donor being spliced to either of the two CAGs^43^. This differential utilization of the tandem splice sites, referred to as NAGNAG splicing, contributes to proteome diversity^43,44^. To test whether the expanded CAG repeats could be acting as alternative acceptor sites, we analyzed the RNA sequencing reads that span the entire repeat tract for relatively short repeats (10 and 22×CAG) that are accessible via short-read sequencing (Supp. Fig. 9A). At these repeat lengths, we observed that the primary acceptor was the first CAG of the repeat unit, accounting for at least 75% of the splicing events for 10×CAG and 50% of the splicing events for 22×CAG (Supp. Fig. 9B-C). It is important to note that PCR amplification, such as during sequencing library preparation, can lead to truncation in repeat number^45,46^. We observe substantial truncations during PCR even when using DNA with a fixed number of CAG repeats (Supp. Fig. 9D). Such truncations increase with repeat number and make it challenging for us to precisely identify the acceptor CAG within the repeat region. Another possibility is that the expanded repeats sequester proteins that alter splice site choice and efficiency. Splicing factors such as MBNL1 and SRSF6 bind RNAs with tandem CAG repeats^6,47,48^. An expanded repeat tract may recruit multiple copies of these splicing factors and affect splice site utilization. Supporting this model, CAG repeat-dependent RNA processing defects have been documented in multiple diseases^6,15,17,18,47^. These two models are not mutually exclusive and future work may shed light on the precise mechanisms driving increased splicing to expanded CAG repeats.

To what extent does splicing to tandem CAG repeats occur in disease? To address this question, we examined the publicly available RNA sequencing datasets generated from post-mortem brains of patients affected by HD, the best studied CAG trinucleotide repeat expansion disease. From this composite dataset (∼8 billion total reads), we mapped only 396 reads adjacent to the *HTT* CAG repeat tract (Supp. Fig. 10A), a notoriously underrepresented region in sequencing data^19^ (Supp. Fig. 10B). We did not find statistically significant evidence supporting splicing to the *HTT* CAG repeat in this dataset beyond the background noise (Supp. Fig. 10A). These observations suggest that splicing to the expanded CAG repeat in *HTT* is likely rare, and occurs at a frequency of <0.5%, barring potential biases in RNA sequencing library preparation. By necessity, these studies were conducted on regions of the brain that were present in the postmortem tissue and may underrepresent the most affected cells. One potential implication of splicing to repeats would be the production of aberrant RAN translation products. RAN translation is only observed in a fraction of cells (up to ∼20% cells in the highly affected striatum, and less frequently in other regions) in HD postmortem tissue^8^. It is possible that splicing to the *HTT* CAG tract is rare and occurs only in a small fraction of cells and is below the detection limit of our current assays. Future studies employing appropriate disease samples and assays with increased coverage in the repeat region may allow us to unequivocally assess the extent of splicing to CAG repeats in disease.

It is also plausible that certain CAG repeat expansion disorders may not involve abnormal splicing to CAG repeats. Most CAG repeat loci in the genome do not serve as splicing acceptors (Supp. Fig. 11A). The presence of tandem CAG repeats, which might act as splice acceptors, may impose constraints on the surrounding sequences, and evolutionary pressure may have purged cis-acting splicing-associated features (such as polypyrimidine tract) from these neighboring regions. To assess whether CAG repeat expansions at new genomic locations (i.e., locations that do not endogenously harbor a CAG repeat) could lead to splicing from adjacent donors, we introduced CAG repeats in an intron of *ACTB* in U-2OS cells (Supp. Fig. 12A). This intron has 8 isolated CAGs, four of which are predicted to be strong splice acceptor sites (3’ MaxEnt 2.76 – 9.47, Supp. Fig. 12B). These native single CAG trinucleotides do not act as splicing acceptors (Supp. Fig. 12C). We incorporated varying numbers of CAG repeats in this intron, isolated monoclonal cells at each repeat number, and characterized the transcribed RNAs using SATCfinder (Supp. Fig. 12D-H). We observed numerous reads where the upstream exon was spliced to the CAG repeats, in a repeat-number dependent manner (Supp. Fig. 12H-I). These reads reflecting splicing to the inserted CAG repeat were ∼1% as abundant as the reads corresponding to the correctly spliced *ACTB* transcript (Supp. Fig. 12J). *ACTB* is an essential gene and only one locus in our polyploid cells was modified with the repeat insertion (see Methods). Although the fraction of transcripts that are mis-spliced due to the repeat insertion is modest, these results underscore that expansions of CAG trinucleotide repeats in an appropriate context can result in de-novo splicing acceptors that sequester adjacent donor sites and produce aberrantly spliced mRNAs.

Our findings add to various molecular mechanisms that contribute to the aberrant proteins that are attributed to RAN translation. Besides CAG repeat-expansion disorders, RAN translation has also been observed in several other repeat-expansion diseases. In fragile X tremor ataxia syndrome (FXTAS), caused by CGG repeat expansion in *FMR1*, the RAN products in the poly-glycine frame results from translation initiation at a near-canonical ACG start site^49^. Likewise, in amyotrophic lateral sclerosis, associated with GGGGCC repeat expansion in *c9orf72*, RAN translation in the glycine-alanine frame is dependent on the presence of a near-cognate CUG initiation codon in the upstream sequence^50^. Another possibility is that the secondary structures formed by GC-rich repeat containing RNAs may provide internal ribosome entry sites facilitating cap-independent translation of the repeats^51–53^. GC-rich repetitive sequences can also result in ribosomal frameshifting, and generate chimeric proteins^50,54–56^. We show that in addition to these mechanisms, aberrant splicing of the repeat-containing transcript followed by canonical or potentially near-canonical translation initiation may also contribute to the production of RAN proteins. The location of repeats within a gene is usually annotated based on the wild-type gene without pathogenic expansion. Splicing aberrations may create new isoforms, and modify exon/intron annotations. Advances in long read sequencing technologies in conjunction with methods that allow mapping repetitive regions may reveal the full-length transcripts that are produced in disease, and help elucidate the disease mechanisms.

In conjunction with prior research, our findings help piece together a model for how CAG repeat-expansions affect RNA processing and sub-cellular localization (see model in Fig. 6). CAG repeats potentiate intermolecular RNA-RNA interactions, and an expanded repeat tract results in the sequestration of the repeat-containing RNA at nuclear foci. These foci co-localize with nuclear speckles and sequester various splicing factors, often inducing missplicing of other transcripts. If the repeat-containing RNA also harbors an appropriate splice donor, this donor may be spliced to the expanded CAG repeats, facilitating its export to the cytoplasm. Splicing may place the repeats in AUG-initiated ORFs. Canonical translation of such repeat-containing ORFs may produce homo-polymeric peptides that form pathogenic aggregates. Our results provide a new lens to examine the role of cis-acting sequences as modifiers of CAG-repeat induced cell toxicity, and suggest that modulating RNA splicing could be a potential target for the development of therapeutic interventions.

## Acknowledgements

We thank David Bartel, Christopher Burge, and members of the Jain lab for helpful discussions. Plasmids containing CAG repeats with flanking sequences from ATXN3, ATXN8, JPH3, DMPK-AS, and HTT were a gift from Laura Ranum. This work was supported by grants from the NIH (R00AG053434), Chan Zuckerberg Initiative, David and Lucile Packard Foundation, and the Smith Family Awards Program. Y.C. was supported by a fellowship from the National Research Foundation of Korea. This paper was typeset with the bioRxiv word template by @Chrelli: www.github.com/chrelli/bioRxiv-word-template

## Author contributions

R.A. and A.J conceived this study, designed experiments, and interpreted results. R.A. performed most experiments and data analysis. M.D. performed immunofluorescence and FISH experiments. Y.C. validated AUG-initiation of CAGRAN by Edman degradation and assisted with ACTB CAG knock-in experiments. K.F. optimized TGIRT reverse transcription conditions. R.A. and A.J. wrote the paper with contributions from all authors.

## Declaration of Interests

The authors declare no competing financial interests. A.J. is a member of the scientific advisory board of Molecular Cell.

## Data availability

The SATCfinder pipeline and full documentation will be available at time of publication. Raw data generated in this manuscript, including sequencing files and unprocessed western blot images, will be available at time of publication.

## Materials and Methods

### Cloning and plasmid generation

Complete sequences for all plasmids used in this study are provided in Supplemental Table 2. Lentiviral transfer plasmids with CAG-repeats and MS2hairpins were previously described^5^, as were the plasmids with 240×CAG repeats with various upstream flanking sequences (including CAG^RAN^ and CAG^FOCI^)^12^. Plasmids with ∼110 CAG repeats and endogenous flanking sequences from *ATXN8, JPH3, DMPK, HTT*, and *ATXN3* were a generous gift from Dr. Laura Ranum^10^. pcDNA3.1(+) EGFP was a gift from Jeremy Wilusz (Addgene plasmid #129020). pCMV-VSV-G was a gift from Bob Weinberg (Addgene plasmid #8454). psPAX2 was a gift from Didier Trono (Addgene plasmid #12260). pX330-U6-Chimeric_BB-CBh-hSpCas9 was a gift from Feng Zhang (Addgene plasmid #42230). Expanded CAG repeats are difficult to amplify using polymerase chain reaction, and make it challenging to incorporate point mutations using site-directed mutagenesis. Instead, mutations were incorporated by using synthetic double stranded DNA fragments that replaced the corresponding fragments in the plasmids. Double stranded DNA fragments with mutations to CAG^RAN^ splice donor and AUG sites (to generate CAG^RAN*donors^ and CAG^RAN*AUGs^, respectively) were prepared by overlap extension PCR (see oligo sequences in Supplemental Table 3), and inserted between MluI and EcoRI sites in CAG^RAN^. CAG^RAN^ constructs with varying repeat lengths were prepared by oligo annealing and inserted between the EcoRI and NotI sites. ATXN8 mutants to AS-AmpR were generated with overlap extension PCR, then inserted between BspHI sites in the ATXN8 or ATXN8^KKQ^ constructs. Splice donors adjacent to the CAG repeat tract were mutated by overlap extension PCR, then Gibson assembled into XbaI and SacI-digested ATXN8^KKQ^. The endogenous 400 bp ATXN8 constructs, and splice donor and AUG mutants, were prepared by overlap extension PCR and inserted between MluI and NotI sites.

All plasmids were propagated in Stbl3 *E. coli* (Invitrogen C737303) grown at 37°C in Luria-Bertani (LB) medium. AS-AmpR mutants grew poorly on LB/agar plates supplemented with 100 μg/mL carbenicillin without an extended outgrowth period (>1 hour). All plasmids were verified by Oxford Nanopore sequencing (Plasmidsaurus or Quintara Bioscience) or by Sanger sequencing using a modified protocol with added 7-deaza dGTP and betaine to allow sequencing through tandem repeats.

### Cell culture

HEK293T (CRL-3216), RPE-1 (CRL-4000), U-2OS (HTB-96), and NIH-3T3 (CRL-1658) were obtained from ATCC and were tested for mycoplasma at least quarterly using MycoAlert (Lonza LT07-318). U-2OS cells stably expressing a TetOn 3G transactivating protein and MS2-hairpin binding protein fused to YFP were previously described^5^. RPE-1 and NIH-3T3 cells expressing the same proteins were prepared similarly by sequential lentiviral transduction and selection. All cell lines were maintained at 37 °C in 5% CO_2_ in Dulbecco’s Modified Eagle Medium (DMEM; Gibco 11965126) supplemented with 10% fetal bovine serum (FBS; Gibco 26140079) and 1% Penicillin-Streptomycin-Glutamine (PSG; Gibco 10378016). Cells were passaged 3 times per week at 1:10 dilution using Dulbecco’s Phosphate Buffered Solution (DPBS; Gibco 14190144) and trypsin (Gibco 25200072). For transient transfections, HEK293T cells were seeded in a 6-well plate at 750,000 cells/well overnight. 2-4 μg of plasmid was mixed with 500 μL Opti-Mem (Gibco 31985070), after which 2.5 μL Plus Reagent and 8 μL Lipofectamine LTX (Invitrogen 15338100) were added. After a 10-minute incubation at room temperature (22 °C), the mixture was added dropwise to wells containing 1 mL Iscove’s Modified Dulbecco’s Medium (IMDM; Gibco 12440053). After 6 hours, the media was exchanged for DMEM with FBS and PSG.

To prepare lentivirus, HEK293T cells were transfected as described above in 6-well plates with 2 μg of transfer plasmid, 0.5 μg envelope plasmid (Addgene #8454), and 1 μg packaging plasmid (Addgene #12260). 24 hours post transfection, the viral supernatant was collected and spun at 24,000×g for 10 minutes at room temperature. U-2OS cells were transduced at varying viral titer in fresh DMEM with FBS and PSG and 10 μg/mL polybrene (Millipore Sigma TR1003G) to increase transduction efficiency. To induce transgene expression, cells were plated to be 80% confluent at time of use and were treated with 1 μg/mL doxycycline (Sigma Aldrich D9891) for 24 hours prior to use. For splicing inhibition, Tet-inducible cells were induced with 1 μg/mL and co-treated with 0.1% DMSO (Sigma Aldrich D2650) or pladienolide B (Santa Cruz Biotechnology, SC391691) at the indicated concentrations (12.5, 25, or 50 nM, depending on the assay) for 24 hours prior to analysis. For splicing inhibition during transient transfection, cells were transfected as above, and after 6 hours, the media was exchanged for DMEM with FBS and PSG, with 25 nM pladienolide B, or 15 μM isoginkgetin (Tocris Bioscience 6483), or 0.1% DMSO as control.

### Western blot

Cells were washed with DPBS and lysed with 160 μL RIPA buffer (25 mM Tris-HCl pH 7.5 (Invitrogen 15567027), 150 mM NaCl (Invitrogen AM9760G), 1% (v/v) NP-40 (Fisher Scientific AAJ19628AP), 1% (w/v) sodium deoxycholate (Sigma-Aldrich D6750), 0.1% (w/v) SDS (Bio-Rad 1610302)) supplemented with 1% (v/v) HALT protease and phosphatase inhibitors (Thermo Scientific 78429) and 125 U/mL Benzonase nuclease (EMD Millipore E1014). The cell lysate was homogenized by passing through a 26-gauge needle (Becton Dickinson 305110) 5 times, and incubated on a nutator for 30 minutes at 4 °C. We noticed that polyglutamine-containing proteins were frequently lost in the pellet fraction after centrifugation at 21000×g or even at 3000×g for 5 minutes. Thus, homogenized lysate was used without further clean-up for electrophoresis. Lysates were incubated with NuPAGE 4× LDS buffer (Invitrogen NP0008) with 50 mM dithiothreitol (DTT; Thermo Scientific R0861) at 70 °C for 5 minutes for most assays. When probing polyalanine-containing proteins, we observed substantial aggregation when lysates were heat denatured at or above 70 °C. Instead, when detecting polyalanine proteins, denaturation was performed at 37 °C for 5 minutes, as is recommended for hydrophobic transmembrane proteins^58,59^, and this denaturation condition substantially reduced polyalanine aggregation (Supp. Fig. 8C). Samples were separated on a Bolt 4-12% Bis-tris polyacrylamide gel (Invitrogen NW04122) run at 200 V for 30 minutes (for 7 kD MW endogenous ATXN8 constructs) or 42 minutes (all other samples). Samples were transferred to PVDF membranes (Invitrogen IB24002) using the iBlot 2 dry blotting system (Invitrogen IB21001). Membranes were blocked in 5% (w/v) skim milk (BD Biosciences 232100) in TBST (tris-buffered saline (Fisher Scientific AAJ60764K3) with 0.1% (v/v) Tween-20 (Fisher Scientific BP337) for one hour at room temperature, then incubated with primary antibodies in 1% (w/v) skim milk in TBST at 4 °C overnight. The membranes were washed 4 times for five minutes each with TBST, then incubated with the appropriate secondary antibody in 1% (w/v) skim milk for one hour at room temperature. After 4 five-minute washes in TBST, chemiluminescence was detected using SuperSignal West Femto Maximum Sensitivity Substrate (Thermo Scientific 34095) with a ChemiDoc XRS+ imager (Bio-Rad). Primary antibodies were used at the following dilutions: mouse anti-FLAG (1:1000, Sigma-Aldrich F1804), mouse anti-HA (1:1000, BioLegend 901501), mouse anti-polyQ (1:2000, Sigma-Aldrich MAB1574), mouse anti-NPT (1:2000, Invitrogen MA5-15275), rabbit anti-β-tubulin (Cell Signaling Tech. 2146). Secondary antibodies: goat anti-rabbit HRP conjugate (Sigma Aldrich A0545), rabbit anti-mouse HRP conjugate (Sigma Aldrich A9044). Band intensities were quantified using ImageJ (version 1.53q)^60^, with background subtraction performed per-lane.

### N-terminal protein sequencing of CAG^RAN^

Four 10-cm dishes expressing CAG^RAN^ fused to BFP at the C-terminus^12^, downstream of the repeat tract, were induced with doxycycline. After 24 hours, the cells were washed once with DPBS and then lysed with 1 mL of RIPA buffer supplemented with HALT protease and phosphatase inhibitors and DTT. The cell lysate was collected by scraping and homogenized by passing through a 22-gauge needle (Becton Dickinson 511055) 10 times, and incubated on a nutator for 30 minutes at 4 °C. The lysate was loaded onto 70 μL of GFP-Trap beads (Chromotek gtma-20) which were prewashed once with RIPA wash buffer (as above, but with 0.15% NP-40). The lysates were incubated on a nutator for 90 minutes at 4 °C, after which the beads were sedimented at 2,500×g for 5 minutes at 4 °C, then washed with 500 μL RIPA wash buffer. After four total wash steps, 80 μL of 2× SDS buffer (Fisher Scientific LC2676) and 8 μL of 1 M DTT was added. The beads were boiled at 95 °C for 10 minutes, then the supernatant was separated on a Bolt 4-12% Bis-tris polyacrylamide gel run at 200 V for 32 minutes, and finally transferred to PVDF membrane using the iBlot 2 dry blotting system. The membrane was rinsed with water, then soaked in 100% methanol (VWR EM-MX0475-1) for 10 seconds, and stained with Coomassie blue staining solution (40% (v/v) methanol, 1g/L Coomassie R250 (Sigma Aldrich 1125530025), 1% acetic acid (Sigma Aldrich 695092) for one minute. The membrane was destained with 50% (v/v) methanol until the destaining solution remained colorless, rinsed with water, and dried at room temperature. Bands extracted from the PVDF membrane were subjected to N-terminal sequencing by Edman degradation (Molecular Structure Facility, UC Davis).

### Generation of *ACTB*^*CAG*^ knock-in cell lines

A CRISPR guide was designed using CHOPCHOP^61^ to select a site which might be able to act as a polypyrimidine tract. We designed a ssDNA repair template with 40×CAG repeats and 40 base homology arms targeting the cut site in intron 3 of *ACTB* (sequence in Supplemental Table 3, and see Supplemental Fig. 12). This repair template was obtained as a synthetic oligonucleotide (IDT) and transfected in HEK293T cells. In brief, 2 μg of plasmid expressing CRISPR-Cas9 and guide (Addgene #42230) and 40 pmol ssDNA repair template were transfected using Lipofectamine 3000 (Invitrogen L3000001) following manufacturers’ instructions, using 7.5 μL of L3000 reagent. A GFP plasmid (0.2 μg) was co-transfected to facilitate isolation of transfected cells. After 48 hours, cells were dissociated using trypsin, resuspended in DPBS, and sorted by fluorescence-activated cell sorting using an Aria III Cell Sorter (BD Biosciences). The top 25% of GFP positive cells were plated as single cells and expanded to monoclonal populations. Approximately 50 clones were isolated and examined for the incorporation of CAG repeats at the desired locus using PCR screening as follows. Approximately ∼5000 cells were pelleted and after removing the residual media, 50 μL DNA QuickExtract solution (Lucigen QE09050) was added. The mixture was vortexed, heated at 65°C for 10 minutes and 95 °C for 5 minutes, and then diluted to 200 μL with nuclease-free water. The locus with expected insertion was amplified by PCR using 5 μL KOD Hot Start 2× Master Mix (Sigma Aldrich 71842-3), 0.2 μL each primer at 10 μM, 0.5 μL DMSO, 1 μL gDNA, and 3.1 μL water, with denaturation at 95 °C for 20 seconds, annealing at 60 °C for 10 seconds, and extending at 70 °C for8 seconds for 40 cycles. The ∼300 bp thus generated amplicons were visualized by 2% agarose gel (Invitrogen 16500500). Unexpectedly, we found wide variability in the amplicon size for 40×CAG reflecting that each clone had a different number of CAG repeats. The number of CAG repeats in each clone were stable and did not measurably vary on our experimental timescales.

We selected 20 clones with variable insertion sizes for further validation using nested PCR. The outer PCR was performed as above, but for 25 cycles with 15 second extension to amplify a ∼1300 bp region. Excess primers were degraded by adding 0.5 μL of thermolabile exonuclease I (NEB m0568) at 37 °C for 15 minutes, followed by enzyme inactivation at 80 °C for 5 minutes. 1 μL of this mixture was used as template in a 25 μL reaction for inner PCR for 25 cycles, amplifying an ∼800 bp region as above. The final amplicons were subjected to PCR clean up (Zymo D4004) according to the manufacturer’s protocol, and single molecule sequencing using Oxford Nanopore (Plasmidsaurus). Reads entirely spanning the CAG repeat insertion site were selected with sequential runs of bbduk with options “k=20 literal={5’ ACTCTCTTCTCTGACCTGAG or 3’ CTCTCTTCTCTGACCTGAGT} rcomp=t hdist=0 mm=f”, then assembled using Canu^62^ v2.1.1 using options “-p asm useGrid=0 genomeSize=1000 minReadLength=100 minOverlapLength=100 corMinCoverage=50 corMaxEvidenceErate=0.15 nanopore-raw {file.fq}”. From 20 clones screened, we found repeats tracts of length 5 – 39×CAG. We speculate that this variability is caused by a hairpin formed by the ssDNA template during repair. The clones we selected for RNA-seq had consistent repeat lengths among the sequencing reads, indicating that a single allele was edited. ∼30% of reads spanning the insert region had a CAG repeat, suggesting the *ACTB* gene is triploid in our HEK293T cells, consistent with prior reports^63^.

### Real-time quantitative and endpoint PCR

RNA and DNA were isolated from ∼10^6^ cells using PureLink RNA (Invitrogen 12183018A) and PureLink Genomic DNA (Invitrogen K182001) mini kits, respectively, according to the manufacturer’s protocols. For reverse transcription, contaminating genomic or plasmid DNA was removed using ezDNase (Invitrogen 11766050), followed by reverse transcription using SuperScript IV VILO master mix (Invitrogen 11766050). Briefly, 1 μg total RNA in a 5 μL volume was incubated with ezDNase at 37 °C for 2 minutes. This reaction was diluted with 3 μL nuclease-free water before addition of 2 μL SuperScript IV VILO master mix. The mixture was incubated at 25 °C for 10 minutes to anneal primers, 50 °C for 20 minutes for reverse transcription, followed by inactivation at 85 °C for 5 minutes.

For quantitative PCR, 2.5 ng of total RNA, or 5 pg of plasmid DNA, or 10 ng of genomic DNA, were quantified with SYBR Green PCR Master Mix (Applied Biosystems 4309155) using the QuantStudio 3 RT-PCR system (Applied Biosystems A28567). For determination of relative abundance of the CAG^RAN^ intron region, primers targeted the intronic region or the 5’ end of the transcript region. Intron abundance was normalized relative to the 5’ end of the transcript. For CAG^RAN*AUG^, which has mutations in the 5’ region, the intron was normalized relative to the 3’ end of the repeats. For estimation of transgene copy number, the 5’ end of the transcript was normalized to primers targeting *ACTB*. See primer sequences in Supplemental Table 3.

For endpoint PCR, 10 ng of total RNA, or 100 ng of genomic DNA was amplified with Advantage GC 2 polymerase (Takara Bio 639114) using 1 M GC Melt and primers spanning the region from upstream of the splice donors to the 3’ end of the repeat. The reactions were cycled 25 (cDNA) or 35 (genomic DNA) times, with denaturation at 94 °C for 10 seconds, annealing at 58 °C for 10 seconds, and extension at 68 °C for 25 seconds. The amplicons were separated on a 3% agarose gel. For sequencing, the PCR reaction was cleaned up with Zymo DNA Clean & Concentrate kit (D4004) and Sanger sequenced (Quintara Biosciences).

### Cell toxicity assays

Cell toxicity assays were performed as previously described ^12^. In brief, ∼10,000 cells were plated into a 6-well plate. The next day, the media was replaced with DMEM with FBS and PSG supplemented with or without doxycycline. After five days, when the cells were approximately 75% confluent, the supernatant containing floating (dead) cells was collected. Cells were washed with DPBS to collect loosely attached cells, and were added to the supernatant. The adherent cells were trypsinized and added to the supernatant. Cells were pelleted at 500×g for 3 minutes, then resuspended in 200 μL DMEM. Dead cells were stained with trypan blue (Invitrogen T10282). The cell count and proportion of dead cells were quantified for ≥ 3 technical replicates using a Countess II FL Automated Cell Counter (ThermoFisher AMQAf1000). The cell count for doxycycline induction was normalized to the without-doxycycline condition for each of three biological replicates per condition. The percent of dead cells reflects the number of cells with non-intact membranes, which take up trypan blue.

### RNA sequencing

Total RNA was isolated from ∼10^6^ (6-well) or ∼8×10^6^ (10 cm dish) cells using the PureLink RNA mini kit. RNA quality was verified by denaturing agarose gel or Bioanalyzer. In-house ribodepletion was performed as described previously^64^. In brief, 2.5 μg of total RNA was mixed with 5 μg of ribodepletion oligos (IDT oligo pool, sequences in Supplemental Table 3) and 2 μL of 5× hybridization buffer (100 mM NaCl, 100 mM Tris-HCl, pH 7.5), to a total volume of 10 μL. The RNA and oligos were hybridized by heating to 95°C for 2 minutes and cooling to 45 °C at −0.1 °C/s. While maintaining samples at 45 °C, 1.5 μL of Hybridase RNase H (Lucigen H39500), 3 μL of 5× RNase H buffer (167 mM NaCl, 50 mM MgCl_2_), and 0.5 μL of water were added, and the reaction was incubated at 45 °C for 30 minutes. The ribodepleted RNA was cleaned using 2× RNAClean XP beads (Beckman Coulter A63987), eluting with 20 μL DNase digestion reaction mixture (1.5 μL Turbo DNase (Invitrogen AM2238), 2 μL Turbo DNase 10X buffer, and 16.5 μL RNase-free water). The residual DNA oligos were digested at 37 °C for 30 minutes, and the resulting mRNA was cleaned using 2× RNAClean XP beads and eluted in 11.5 μL nuclease-free water.

The ribodepleted RNA (typically ∼5% of input, or about 100 ng) was reverse transcribed using a group II intron reverse transcriptase (TGIRT, Ingex) using primers provided by the xGen library preparation kit (IDT 10009813). In brief, 10 μL ribodepleted RNA was mixed with 4 μL 5× TGIRT buffer (2.25 M NaCl, 25 mM MgCl_2_, 100 mM Tris-HCl, pH 7.5; Ambion RNase-free Buffer Kit AM9010), and 1 μL random hexamer primers (xGen). The mixture was heated to 94 °C for 12 minutes to induce RNA fragmentation. After cooling at 4 °C for 2 minutes to allow primer binding, 1 μL 100 mM DTT (xGen), 1 μL TGIRT enzyme, and 1 μL RNase inhibitor (xGen) was added and incubated at room temperature for 10 minutes to allow TGIRT binding to RNA-primer duplexes. To initiate reverse transcription, 2 μL dNTPs (xGen) were added, followed by sequential incubation for 15 minutes each at 20 °C, 42 °C, 55 °C, and 65 °C for elongation. This step-wise protocol is needed for TGIRT to extend the unstable RNA-DNA duplexes formed by random hexamer primers^65^. At completion, TGIRT was dissociated from the RNA-DNA duplexes by addition of 1 μL 5 M NaOH (VWR BDH7225-1) with incubation at 95 °C for 3 minutes, then neutralized with 1 μL 5 M HCl (VWR BDH7419-1). The remainder of the library preparation was performed according to the xGen protocol except that post-ligation, the library was eluted in 30 μL nuclease-free water, followed by a modified protocol for final amplification and incorporation of sequencing adaptors. The final amplification was performed in a 50 μL reaction with 1 μL Advantage GC2 polymerase, 10 μL 5× GC2 buffer, 5 μL (1M) GC melt, 1 μL of 10 mM dNTPs (NEB N0447L), 1 μL GC2 polymerase, 4 μL xGen adapter mix, and 29 μL eluted library. Typically, ∼10 cycles of PCR with denaturation at 94 °C for 20 seconds, annealing at 58 °C for 20 seconds, and extension at 68 °C for 45 seconds, for final amplification yielded sufficient material for quality control and sequencing.

### SATCfinder pipeline and RNA-seq analysis

The SATCfinder pipeline and full documentation will be made available at time of publication. In brief, the SATCfinder pipeline first selected reads containing ≥3×CAG or CTG, along with their read mates, if available, using bbduk (BBTools 38.86)^66^ with options “k=9 hdist=0 mm=f literal=CAGCAGCAG rcomp=t”. At this step, read pairs not containing any repeats were discarded. Low quality read ends and sequencing adapters were removed from the reads using cutadapt (version 3.7)^67^ with commands “-a AGATCGGAAGAG --error-rate=0.1 --times=1 --overlap=5 --minimum-length=20 -quality-cutoff=20”. Next, the CAG-selected reads were processed using a custom Python script, fastqToTrimmedSAM.py. This script converts FASTQ files to unmapped SAM files, during which CAG/CTG repeats (minimum 3, maximum unlimited) are computationally trimmed from the read according to the following rules: (1) CAG repeats and anything downstream of the repeat, is trimmed and saved to a SAM field; (2) CTG repeats, and anything upstream of them, is trimmed and saved to a SAM field. In both cases, the length of the trimmed repeat tract (but not any additional removed sequences) is saved in a SAM field. Repeats will be trimmed from both reads if paired end reads are provided. Reads with at least 15 bases post-trimming and their corresponding read mates are stored in the SAM file. These reads were then aligned to hg38 with GRCh GTF annotation file (38.93) using the short-read alignment tool STAR (version 2.7.1a)^68^. STAR alignment arguments were “--outSAMtype BAM SortedByCoordinate --outSAMunmapped Within --genomeFastaFiles {construct.fa} --readFilesType SAM {library type}”, where construct.fa refers to a relevant transgene fasta sequence if required, and library type refers to SE or PE for single and paired end libraries respectively.

The aligned BAM files were indexed using samtools (version 1.1)^69^. Trimmed reads were separated from their read mates and stored in a BAM file which only contains reads with trimmed repeats. This file is then processed by a second python script, findCAGends.py. This script takes as input a BAM file, along with genomic coordinates and strand information for a region of interest, and uses pysam (version 0.16.0.1) to generate a .csv file containing the base position and the number of CAG ends at that base for the given region. This csv file can then be used in standard plotting software to generate bar plots as in Fig. 1A-B, which were prepared with Prism. The percentage of CAG ends reflecting splicing to a CAG repeat was calculated using the number of CAG ends more than 10 bases from the repeat divided by the total number of CAG ends. To estimate the percent of AUG-initiated transcripts after splicing, we examined each splice junction supported by more than 100 reads and asked if that junction would generate an AUG-initiated ORF or not. If yes, we counted the % of CAG ends that defined that junction as AUG-initiated. To find the number of CAG removed during splicing, we examined trimmed and mapped reads that aligned to a region upstream of the repeat tract (either at a splice donor, or directly adjacent to the repeat), that also contained at least 10 bases from the 3’ end of the repeat in their trimmed sequences.

For standard (non-trimmed) analysis, RNA-seq libraries were processed through cutadapt as previously described and then aligned to hg38 using STAR with arguments “--outSAMtype BAM SortedByCoordinate -- outSAMunmapped Within”, along with “--genomeFastaFiles {construct.fa}” if a transgene was expressed. Transcript assembly was performed with StringTie (2.2.1)^29^ with default options, using all reads mapping to the construct.

Relative expression for SV40, CMV, and the cryptic colE1 promoter was calculated by the mean read counts across three replicate libraries for (1) the SV40 transcript, as detected by StringTie; (2) the region between the CMV promoter and CAG repeats, as the 3’ end of the CAG repeat tract could be present on both initiated by CMV and by colE1; and (3) the region of the colE1-initiated transcript detected by StringTie without splicing to the CAG repeat tract.

Reads mapping to genes were quantified by featureCounts (version 1.6.2)^70^ using arguments “-g gene_name -f -O”. For reads mapping to exons, we first flattened the GTF (flattenGTF), then used featureCounts with additional arguments “--F SAF -t exon –largestOverlap”.

For HD patient RNA-seq analyses, we utilized the following datasets: GSE64810, GSE124664, GSE79666, GSE129473, GSE109534, GSE159940,

GSE144559. These datasets comprise paired end sequencing (≥ 75 bp reads) from HD patients and unaffected patients. We included only HD-affected patients in our analysis. Of note, the dataset GSE71191 was released only as a composite dataset of reads mapping to the HTT region from multiple patients, and thus we likely overestimate the prevalence of reads mapping to the HTT exon 1 region. These data were analyzed using SATCfinder as previously described.

### Splice site analysis

Sashimi plots without splice junction arcs, as in Fig. 1A-B, were generated with ggsashimi(version 1.1.5)^71^ with arguments “--out-format pdf --mincoverage 100000 --shrink --fix-y-scale”. To find CAG ends, we used a custom Python script, findCAGends.py as described above.

MaxEnt scores for 5’ and 3’ splice sites were calculated using the MaxEntScan web-server (http://hollywood.mit.edu/burge-lab/maxent/Xmaxentscan_scoreseq.html,^25^), or using maxentpy (https://github.com/kepbod/maxentpy/), a python wrapper for the same model. Genome-wide analysis was performed using a custom python script which calculated the 5’ MaxEnt score for every annotated splice donor site in hg38 using the previously described GTF file. No corrections were made for the ∼1% of introns which are processed by the minor spliceosome.

To locate repeats in the genome, and to find splice acceptors with tandem CAG repeats, we used a custom python script, findRepeatsInGenome.py, which generates a csv file with repeats assigned to genomic regions (gene, intron, exon, or intergene). To find CAG repeat acceptors, we calculated the distance in bases between the annotated exon junctions and the left-most coordinate of CAG repeats within each exon. A distance less than zero meant that the CAG repeats at least partially overlapped with the annotated exon junction and thus defined as the splice acceptor site. For example, …YYYYCAGCAGCAGCAG… (where Y is the polypyrimidine tract and the exonic bases are indicated in bold) would have a distance of −3, because the CAG repeat begins 3 bases upstream from the exon boundary. A distance greater than zero indicated a small intervening sequence between the annotated acceptor and pure CAG repeat. Acceptors with interruptions between the acceptor ‘CAG’ and pure CAG tract were not included in our analysis, but are included in Supplemental Table 2 as a reference.

### Fluorescence microscopy

For live-cell imaging with the MS2 system, ∼10,000 cells were plated on glass bottom 96-well plates (Brooks, MGB096-1-2-LG-L) and induced with doxycycline (and any other treatments, as indicated) the following day. Cells were imaged 24 hours after induction with a Dragonfly 505 spinningdisk confocal microscope (Andor Technologies) equipped with a piezo Zstage (ASI) and an iXon Ultra 888 EMCCD camera. Pin-hole size was kept at 40 μm. Z-stacks were acquired with a step size of 0.3 to 0.5 μm. Live cells were imaged in a humidified chamber (OKO labs) maintained at 37 °C and 5% (v/v) CO_2_ using a 100× oil immersion objective NA 1.45 (Nikon MRD01905, pixel size 121 nm × 121 nm). For quantification of cells with aggregates, cells were imaged with a 20× air objective NA 0.75 (Nikon MRD00205). Fixed cells were imaged as above, but at room temperature. DAPI was excited with a 405-nm laser, and fluorescence was collected using a 445/46 bandpass filter. YFP was imaged using a 488-nm laser and corresponding 521/38-nm band pass emission filter. Cy5 and Atto647Nlabeled samples were imaged using 640-nm laser line and a 698/77 bandpass emission filter.

### RNA FISH

Cells were fixed with a solution of 75% (v/v) methanol and 25% (v/v) acetic acid for 10 min at 4 °C. Fixed cells were washed three times with a wash solution (nuclease-free water with 300 mM NaCl, 30 mM sodium citrate (Calbiochem 567446), 10% (v/v) formamide (Sigma Aldrich 47671), and 0.1% (v/v) NP-40 substitute) at room temperature. Hybridization was performed at 37 °C for 3 hours with 200 nM Atto647N-conjugated probes targeting the CAG repeats. The probes were dissolved in hybridization buffer (100 mg/mL dextran sulfate (Sigma Aldrich D8906), 10% (v/v) formamide, 300 mM NaCl, and 30 mM sodium citrate in nuclease-free water). After hybridization, cells were washed for 30 min with wash solution and counterstained for 30 min with wash solution containing 0.5 μg/mL DAPI (Sigma Aldrich D9542); both this wash step and DAPI staining were performed at 37 °C. Cells were then washed three times with phosphate buffered saline (PBS; Gibco 20012-027), and kept in PBS for imaging.

### Immunofluorescence

Cells were fixed with 2% (v/v) formaldehyde (Fisher Scientific 28906) in PBS for 40 min, washed four times with PBS, and then permeabilized with 0.1% (v/v) Triton-X-100 (Sigma Aldrich T8787) in PBS for 10 min at room temperature. Cells were then blocked in 0.45-μm filtered 3% (w/v) bovine serum albumin (BSA; Sigma-Aldrich A7906) in PBS. Primary antibodies targeting poly-glutamine (Sigma-Aldrich MAB1574) were diluted 1:100 in 1% (w/v) BSA in PBS and incubated with cells for 1 h at room temperature. Cells were washed three times with PBS and then incubated with Cy5-conjugated donkey anti-mouse (Jackson Immuno Research Labs 715175151) secondary antibody at 1:2,000 dilution in 1% (w/v) BSA in PBS for 1 h at room temperature. After three washes with PBS, cells were counterstained with DAPI solution (PBS containing 0.5 μg/mL DAPI) for 3 min, washed three times with PBS again, and then kept in PBS for imaging.

### Quantification and statistical analyses

Toxicity assays, quantitative PCR, cells with aggregates, and nuclear/cytoplasmic RNA ratios were analyzed by two-tailed unpaired Student’s t-test. The percentage of endogenous transcripts reflecting splicing to annotated CAG acceptors was analyzed by a Fisher’s exact test. Statistical tests are further described in figure legends.

**Supplemental Figure S1.**
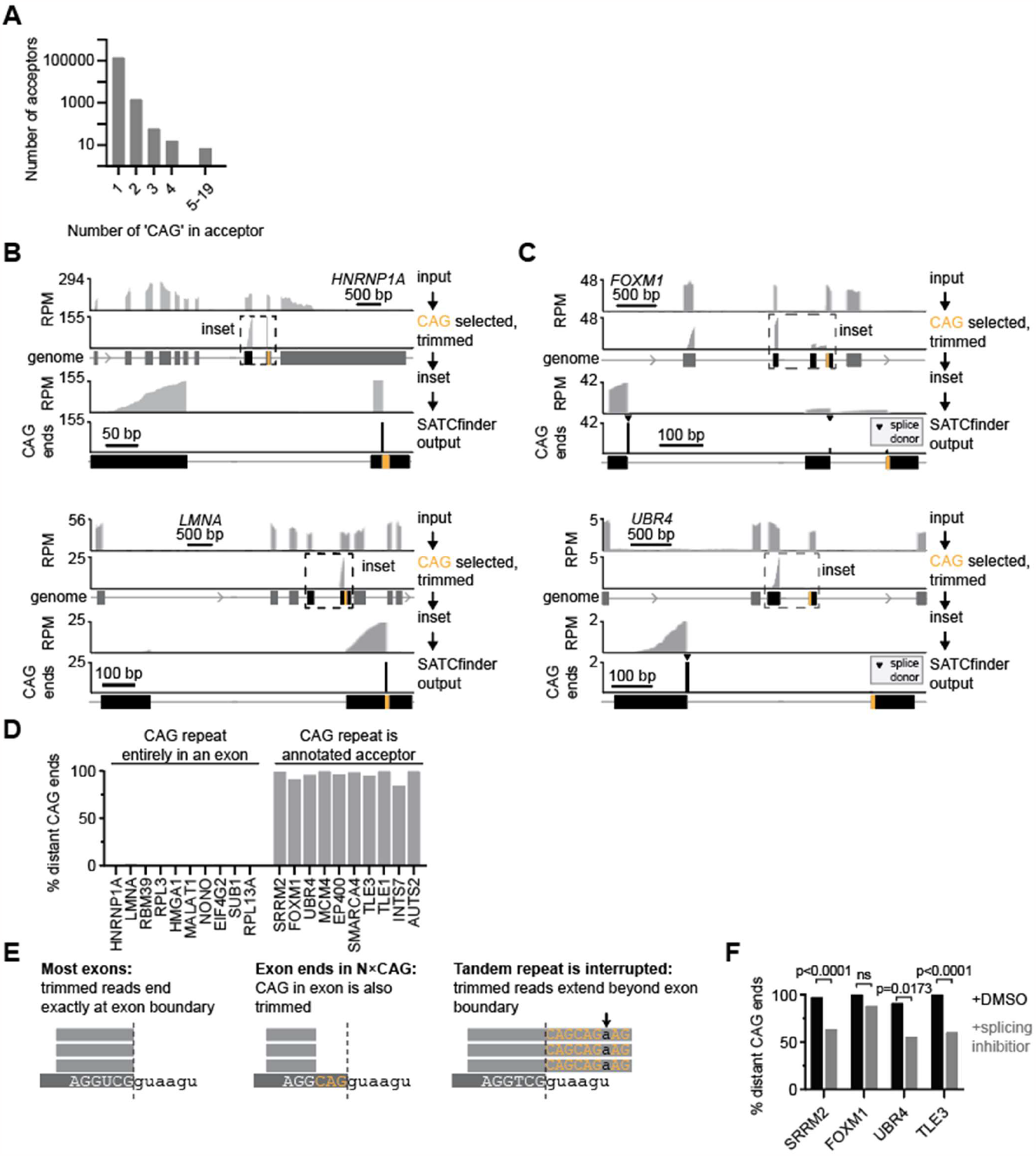
Validation of the SATCfinder pipeline. A. Number of annotated splice acceptor sites in the human genome where one or more CAG triplet occurs at the exon boundary. **B, C**. Additional representative genes comparing standard RNA sequencing analysis to SATCfinder output. **D**. Percentage of CAG ends mapping distal from the CAG repeat for genes without and with annotated CAG repeat acceptors. **E**. Schematic for the various scenarios for the alignment of trimmed reads when CAG repeats act as splicing acceptor sites. In the most cases, the CAG end coincides with the last base of the exon. In some cases, the splice donor may have a **CAG**GU motif, as in 22.8% of human donors. In such cases, the terminal CAG(s) from the 5’ splice site will also be trimmed by our pipeline. Lastly, if the acceptor CAG repeat has interruptions, the CAG end may extend beyond the splice donor. These bases will fail to map and will typically be soft-clipped by the alignment software. **F**. Percentage of CAG ends reflecting splicing to the annotated tandem CAG 3’ splice site in the presence of splicing inhibitor 25 nM pladienolide B, or 0.1% DMSO as control. Significance values in F are calculated using Fisher’s exact test.

**Supplemental Figure S2.**
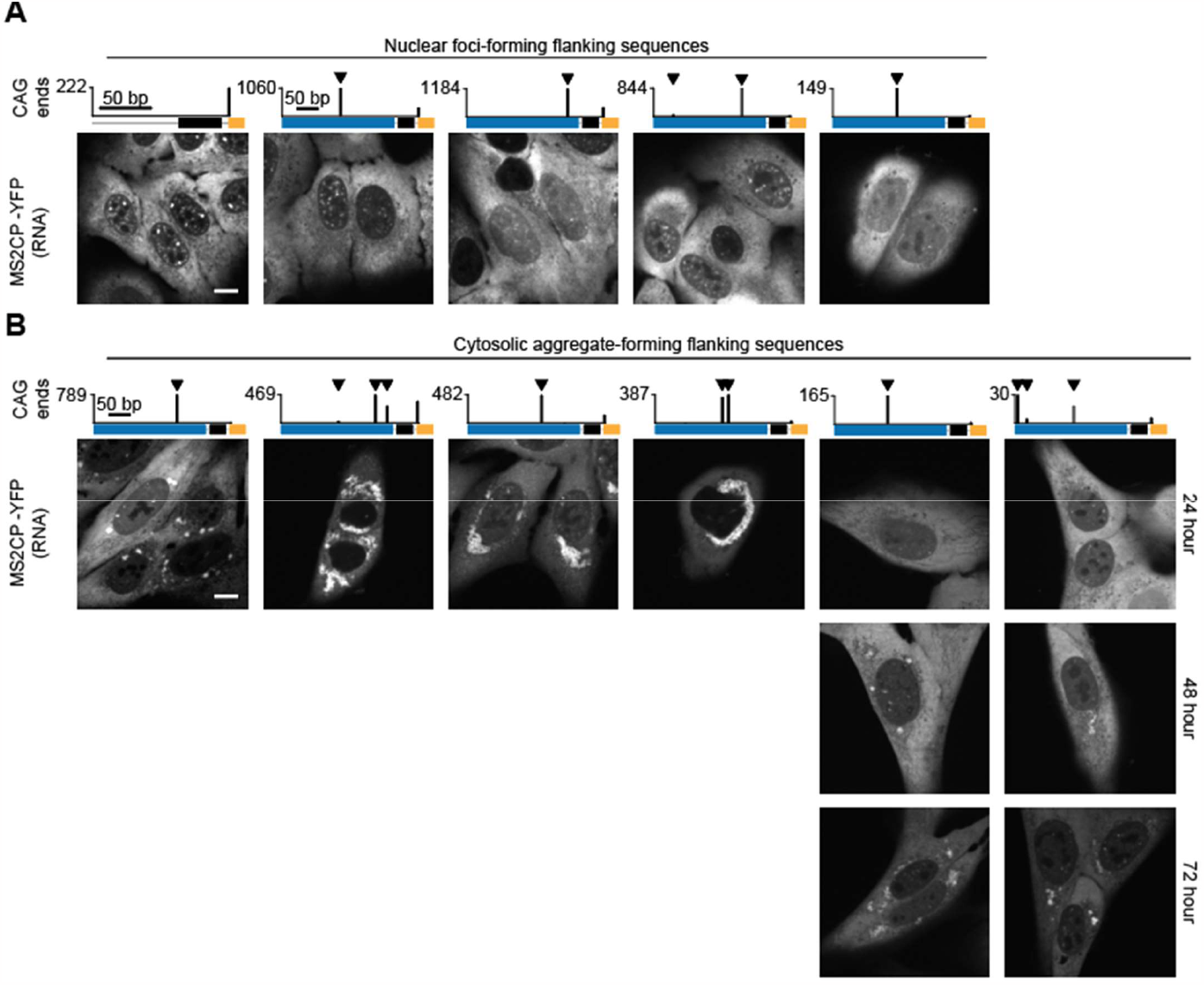
A, B. Characterization of RNA splicing and sub-cellular localization in our library with various 250 base flanking sequences cloned upstream of 240×CAG repeats that exhibit nuclear foci (A) or RAN translation and cytoplasmic aggregates (B)^12^. *Top*, SATCfinder sequencing output for the indicated CAG constructs, in addition to representative examples in Figure 2. *Bottom*, Representative fluorescent images of cells expressing the indicated constructs induced for 24 hours. Two cytosolic-aggregate forming constructs that initially exhibit smaller aggregates are also shown at 48- and 72-hour post induction. Micrographs are representative of ≥ 2 independent experiments. Scale bar, 10 μm.

**Supplemental Figure S3.**
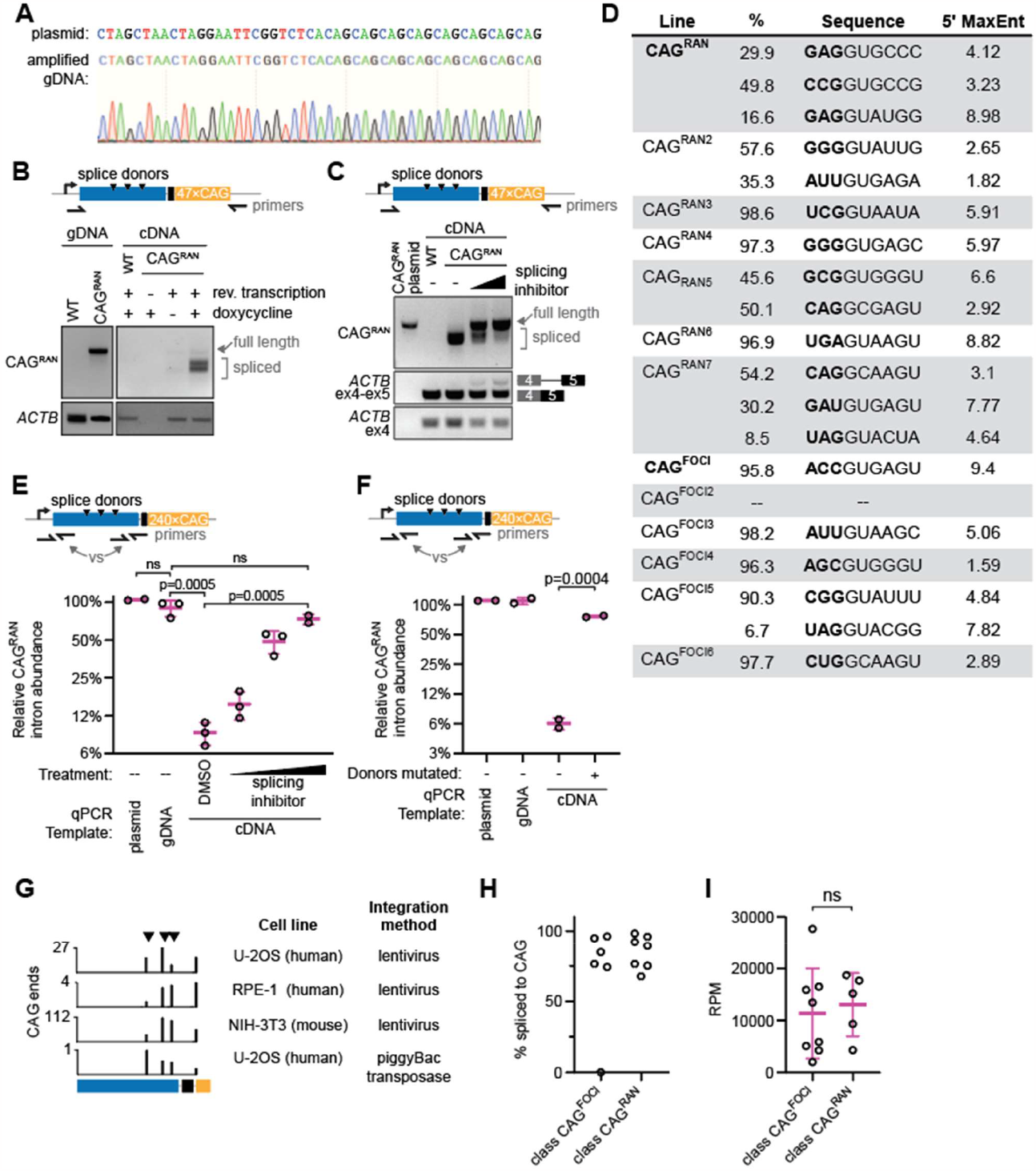
Validation and quantification of splicing to CAG repeat tract. A. Sanger sequencing trace for representative CAG^RAN^ construct amplified from genomic DNA, showing that the desired sequence is integrated into the genome. **B, C**. End-point PCR spanning the CAG repeat for CAG^RAN^ flanking sequence for genomic DNA, plasmid DNA, or cDNA in the presence or absence of induction of CAG^RAN^ expression by 1 μg/mL doxycycline. Schematic depicts the location of primers and observed 5’ splice sites. Splicing to CAG repeats results in a spliced RNA that is smaller than the genomic insert. Primers targeting a region in an exon of *ACTB* are used as control, or spanning *ACTB* exon4-5 junction control for splicing inhibition by 25 or 50 nM pladienolide B in panel C. **D**. Sequences of the identified splicing donors in the CAG flanking sequences library with corresponding percentage of reads arising from each donor. The remaining CAG-containing reads arise from transcripts where the CAG repeat tract does not serve as splicing acceptor. CAG^FOCI2^ does not have a substantial upstream flanking sequence and no splicing to CAG repeat tract is observed. **E, F**. Real-time quantitative PCR for CAG^RAN^ or CAG^RAN*donors^ intron region normalized to the expression of the 5’ end of the CAG^RAN^ transcript. For cDNA, CAG^RAN^ expression was induced in the presence of splicing inhibitor (pladienolide B, at 12.5, 25, or 50 nM), or 0.1% DMSO (-) as control. **G**. SATCfinder sequencing output for the representative CAG^RAN^ flanking sequence transduced into various human and mouse cell lines by lentivirus or by the piggyBac transposase. CAG^RAN^ expression was induced in the presence of splicing inhibitor (25 or 50 nM pladienolide B), or 0.1% DMSO (-) as control. **H**. Percentage of CAG ends reflecting splicing to the CAG repeat for constructs that produce RNA foci only or exhibit RAN translation. Each data point is one construct. **I**. Normalized expression of constructs. Each data point is one construct. Data for C, G, and H show the mean ± SD for at least two independent RNA or DNA isolations, where each data point is an independent experiment. Significance values in E, F, and I are calculated using Student’s t-test. End-point PCR experiments and Sanger sequencing are representative of ≥ 2 independent experiments.

**Supplemental Figure S4.**
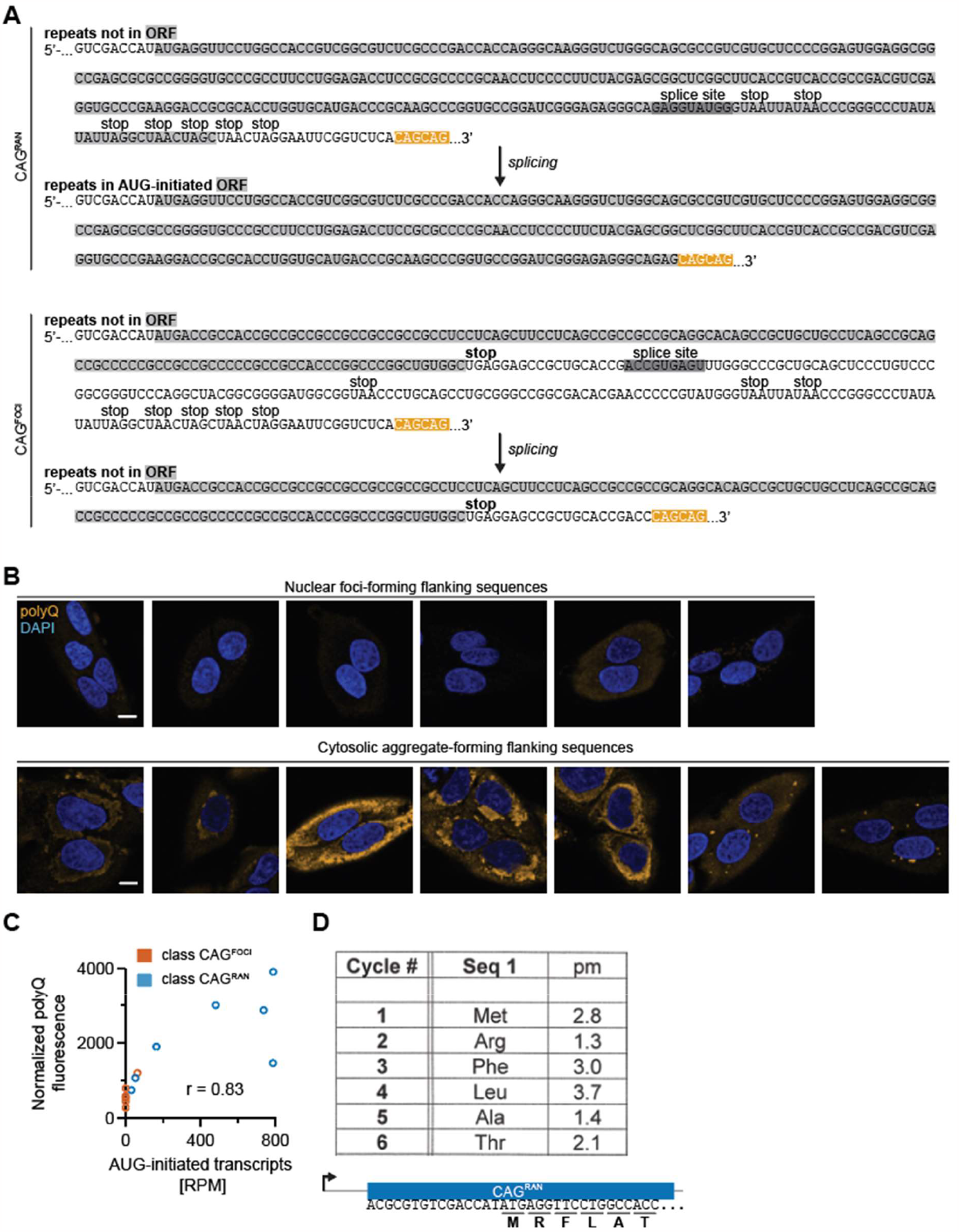
Polyglutamine-containing proteins in the CAG flanking sequence library result from splicing to the CAG repeat. A. Full sequence schematic for splicing events in the representative CAG^FOCI^ (top) and CAG^RAN^ (bottom) flanking sequences. In the absence of splicing, stop codons prevents translation of the repeats regardless of upstream ORFs. After splicing, the repeat tract may be embedded in an AUG-initiated ORF. **B**. Representative immunofluorescence micrographs staining for polyQ-containing proteins in the various construct that either yield RNA foci (top) or cytoplasmic aggregates (bottom). Cell nuclei are stained with DAPI. **C**. Correlation of mean polyQ fluorescence normalized to cell area, in relation to the percentage of repeat-containing transcripts where the repeats are observed in AUG-initiated ORFs. Each data point represents one construct. **D**. Results from N-terminal sequencing of the CAG^RAN^ construct by Edman degradation. A portion of the CAG^RAN^ sequence and the translation of that sequence is indicated. Micrographs and quantification in B and C are representative of ≥ 2 independent experiments with ≥ 15 cells.

**Supplemental Figure S5.**
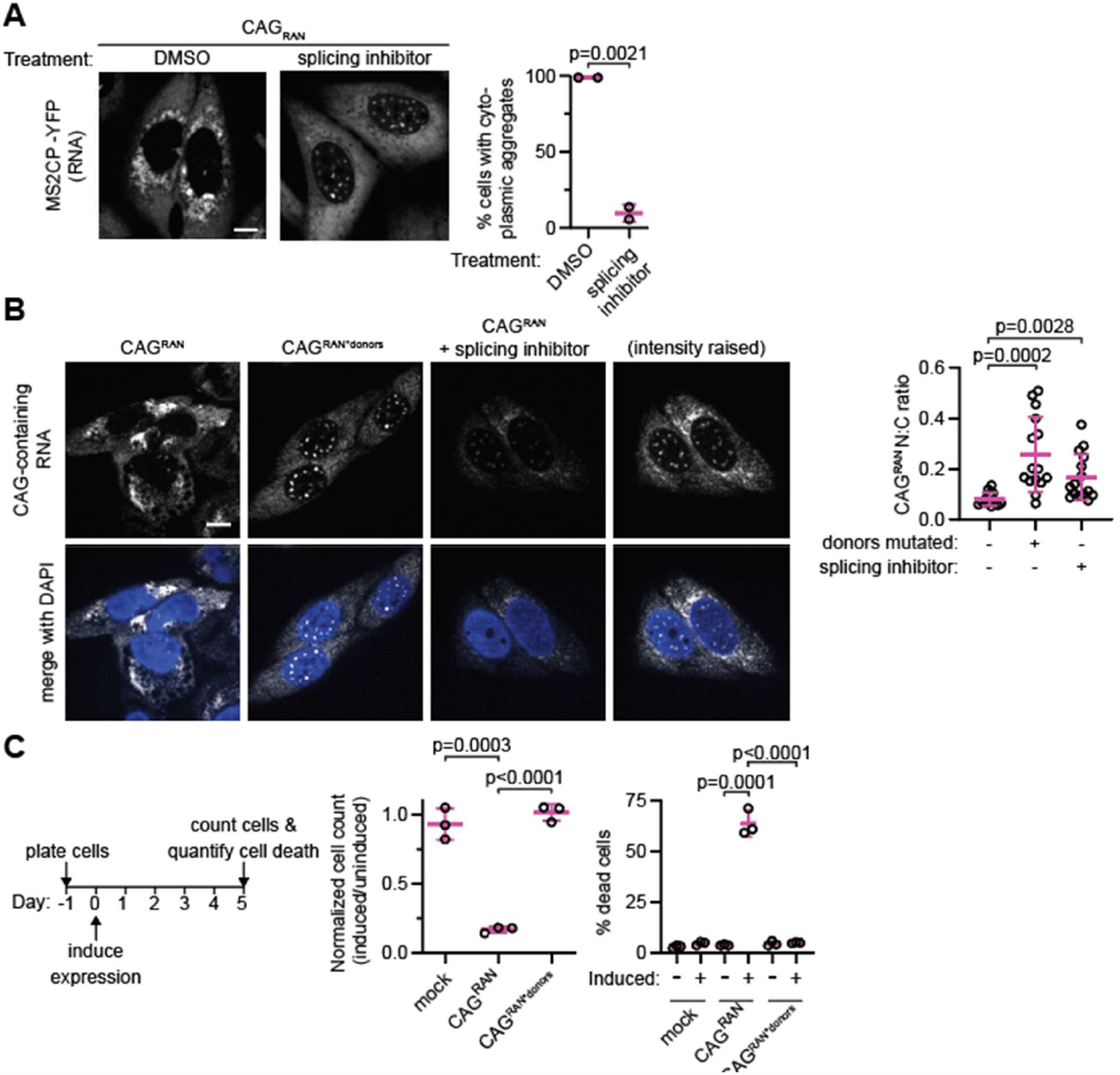
Splicing and ORFs dictate the localization of CAG-repeat containing RNAs. A. *Left*, Representative fluorescent images of cells expressing CAG^RAN^ in the presence of 25 nM pladienolide B (splicing inhibitor) or 0.1% DMSO (DMSO). *Right*, quantification of the % of cells with cytoplasmic RNA aggregates. Each data point represents ≥ 500 cells quantified in an independent experiment. **B**. *Left*, Representative micrographs of cells expressing CAG^RAN^ or CAG^RAN^ with mutated splice donors (CAG^RAN*donors^), detected by fluorescence in-situ hybridization with a 7×CTG probe. Cells were co-treated with 0.1% DMSO, or with 25 nM pladienolide B (splicing inhibitor). Cell nuclei are stained with DAPI. *Right*, Quantification of the nuclear to cytoplasmic ratio for CAG-repeat-containing RNAs. Each data point is an individual cell, n = 15. **C**. Quantification of cell death from expression of CAG^RAN^ or CAG^RAN*donors^. Each data point is a separate biological replicate. Data were normalized first to the uninduced condition, then to the mock parent cell line without a repeat-containing construct. Significance values in A-C are calculated using Student’s t-test. Data in A-C are summarized as mean ± SD. Scale bar, 10 μm.

**Supplemental Figure S6.**
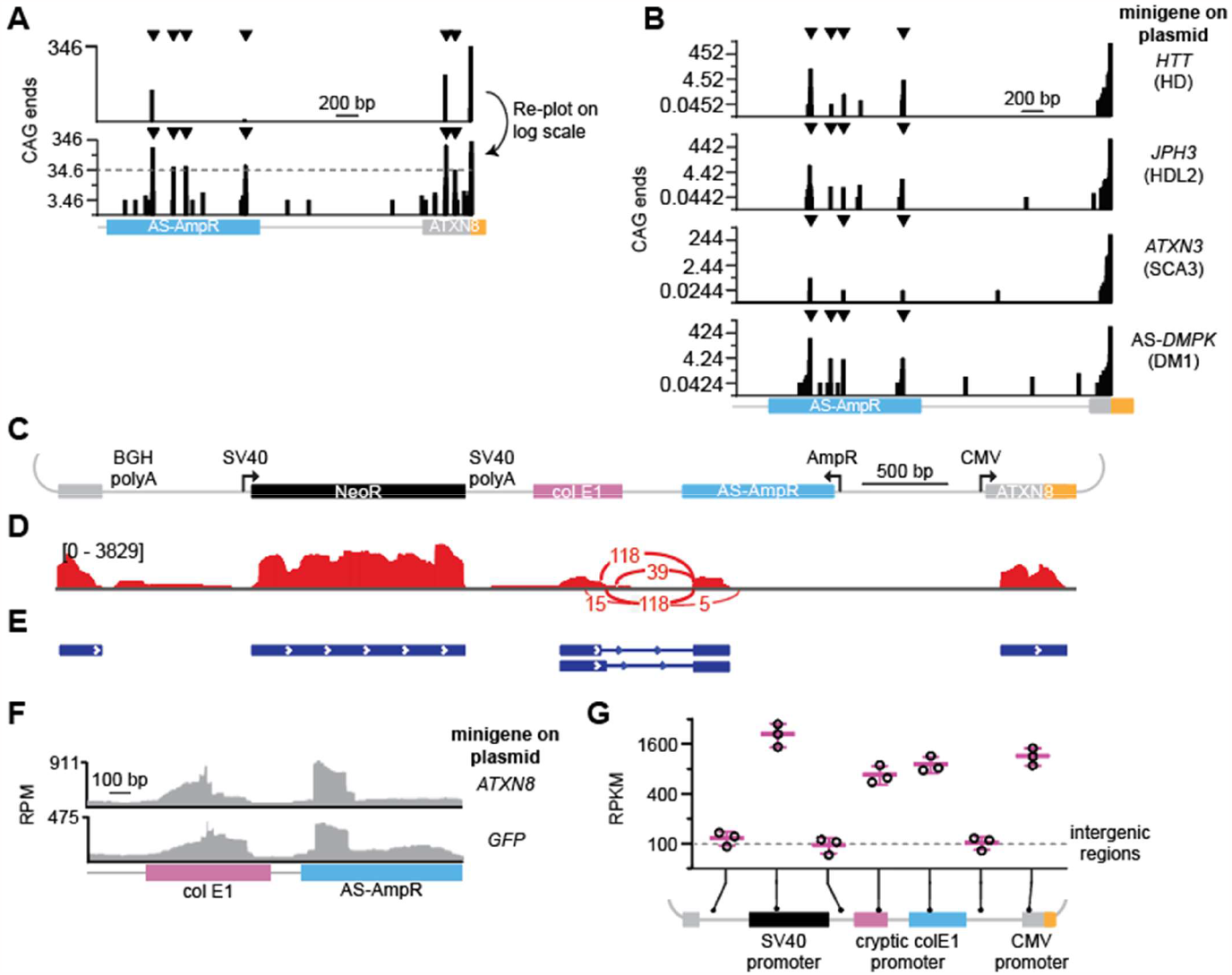
Expanded CAG repeats with native flanking sequences from the disease-associated genes act as splice acceptors. A, B. SATCfinder output for the indicated constructs expressing ∼100×CAG repeats flanked by endogenous sequences from the respective disease-causing gene, with the y-axis on log scale to emphasize the spectrum of 5’ splice sites. **C**. To-scale map for the ATXN8 plasmid with the promoter directionality marked by an arrow and the polyadenylation sites (polyA) indicated. **D**. Sashimi plot for the ATXN8 construct. The y-axis represents the number of reads covering each base. Splice junctions are indicated as arcs between donor and acceptor, with the number of reads spanning the junction overlaid on the arc. No junctions to CAG are depicted because the CAG repeats are trimmed. **E**. Transcript isoforms predicted by StringTie. **F**. RNA sequencing depth of coverage for the indicated region, where cells were expressing either ATXN8 or GFP from the CMV promoter. **G**. Comparison of the abundance of transcripts expressed from the indicated promoters. Each data point represents a separate transfection.

**Supplemental Figure S7.**
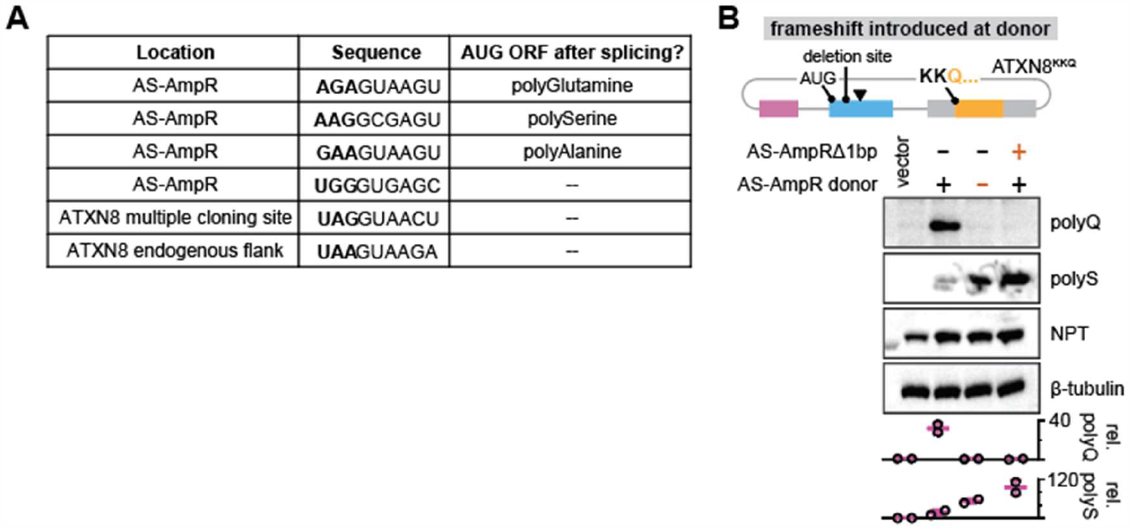
Splicing to CAG repeats in ATXN8 mini-gene results in repeat-containing ORFs that account for aberrant translation products. A. Location and AUG-initiated ORFs generated upon splicing in the ATXN8 construct. **B**. Immunoblots from cells expressing the indicated ATXN8^KKQ^-derived constructs. Band intensities are normalized first to NPT (neomycin phosphotransferase), expressed in *cis* from the plasmid, and then to the control expressing GFP (vector). Tubulin is included to show equivalent loading between conditions, but is not used for normalization due to potential variations in transfection efficiency. Immunoblots and quantification of relative abundance (as mean ± SD) are representative of ≥ 2 independent transfections.

**Supplemental Figure S8.**
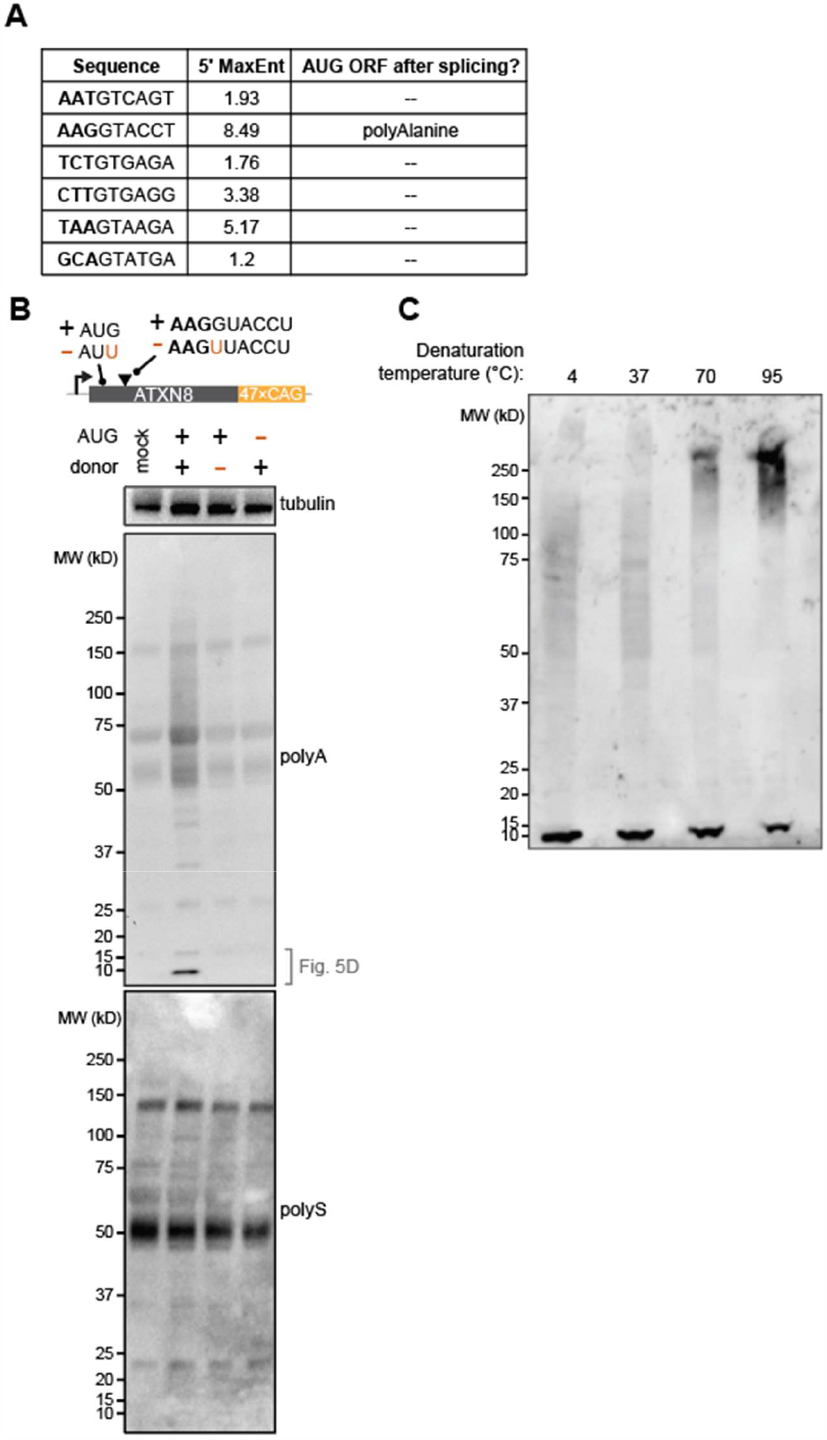
Native upstream flanking sequence in ATXN8 provides a splicing donor and splicing to CAG repeats results in an ORF with the CAG repeats in the polyalanine frame. A. Sequences of the identified 5’ splice sites in the *ATXN8* upstream sequence (400 bp), along with AUG-initiated ORFs resulting from splicing. **B**. Immunoblot for the indicated samples where a HA epitope is in the polyalanine frame and a FLAG epitope is in the polyserine frame. Polyalanine blot shows the full membrane for Fig. 5D with high-MW smear, which we believe reflects aggregated species. Proteins consisting mostly of alanine have previously been shown to run as an extended smear^10^. **C**. Related to panel B, representative blot for the same samples of polyalanine-containing protein denatured at 4 °C, 37 °C, 70 °C, or 90 °C for 5 minutes. With increasing heat, the polyalanine high-MW smear becomes more abundant. This membrane was stripped and re-probed for HA in the polyalanine frame. Immunoblots are representative of ≥ 2 independent experiments.

**Supplemental Figure S9.**
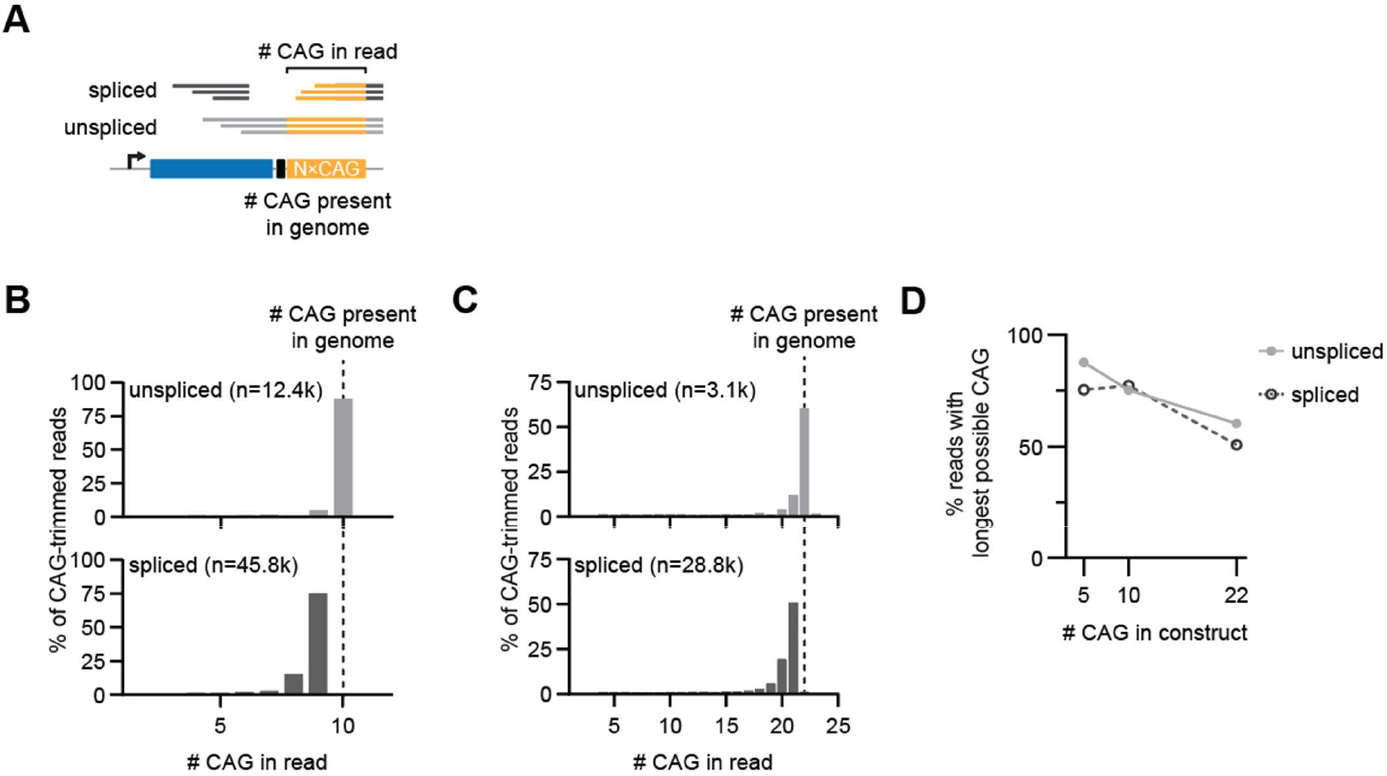
The first CAG triplet of the repeat tract is the primary splice acceptor. A. Schematic for data analysis. The number of CAG repeats was extracted from reads mapping to both the 5’ and 3’ end of the repeat. **B, C**. Histogram for the number of repeats present in RNA-seq reads. Comparison with the number of repeats present in the genome allows estimation of how many CAG triplets are removed during splicing. **D**. Quantification of repeat truncation in PCR during library preparation, which obfuscate the quantification of the number of CAG triplets that are removed due to splicing. If the first CAG triplet of N×CAG acts as the 3’ splice site, an unspliced read with the longest CAG tract would have N×CAG and a spliced read would have N-1×CAG.

**Supplemental Figure S10.**
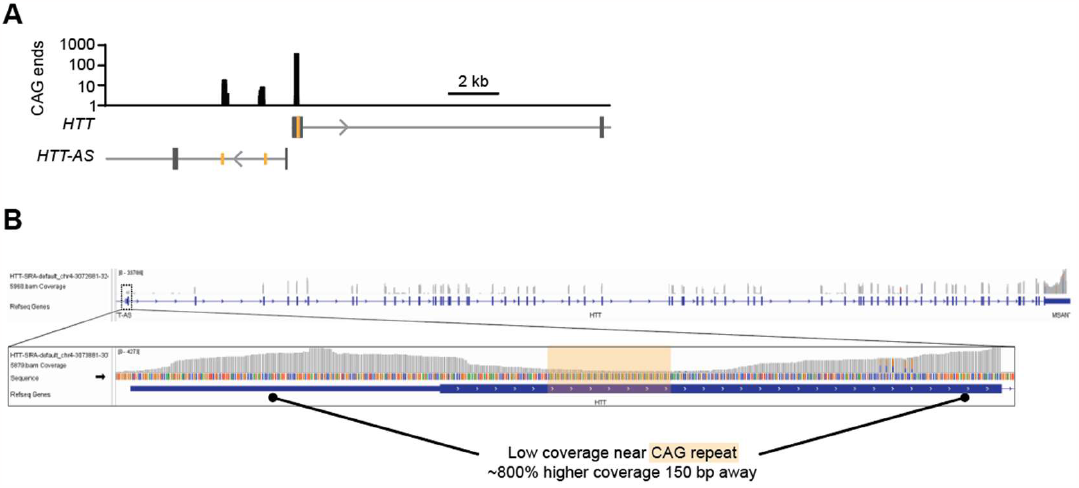
Analysis of publicly available HD patient brain RNA-sequencing datasets. A. SATCfinder output for a 20-kb region centered on *HTT* exon 1. **B**. Raw IGV output for HTT exon 1 for the same datasets as in panel A, but mapped through a standard RNA-sequencing pipeline showing reduced coverage in the regions adjacent to the CAG repeats.

**Supplemental Figure S11.**
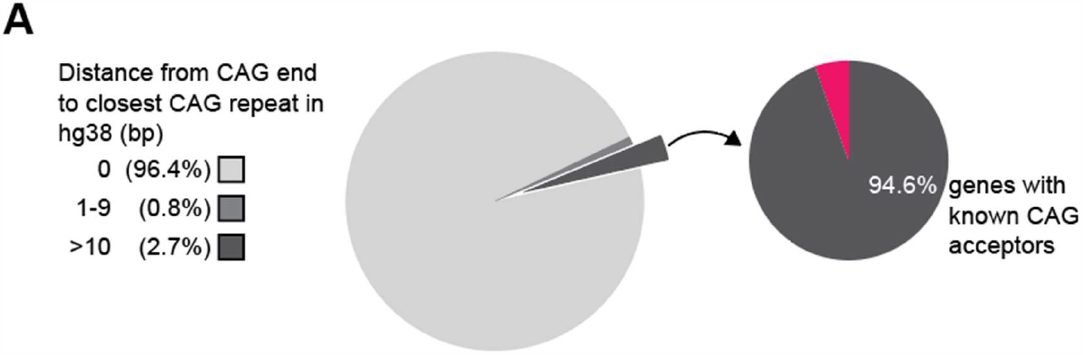
Most CAG repeat sites in the genome do not act as 3’ splice sites suggesting selection pressure on the flanking sequences. A. Quantification of the location of CAG ends in RNA compared to their corresponding genomic locus. Most CAG ends in RNA map to the corresponding location in the genome, suggesting that they do not function as splicing acceptors.

**Supplemental Figure S12.**
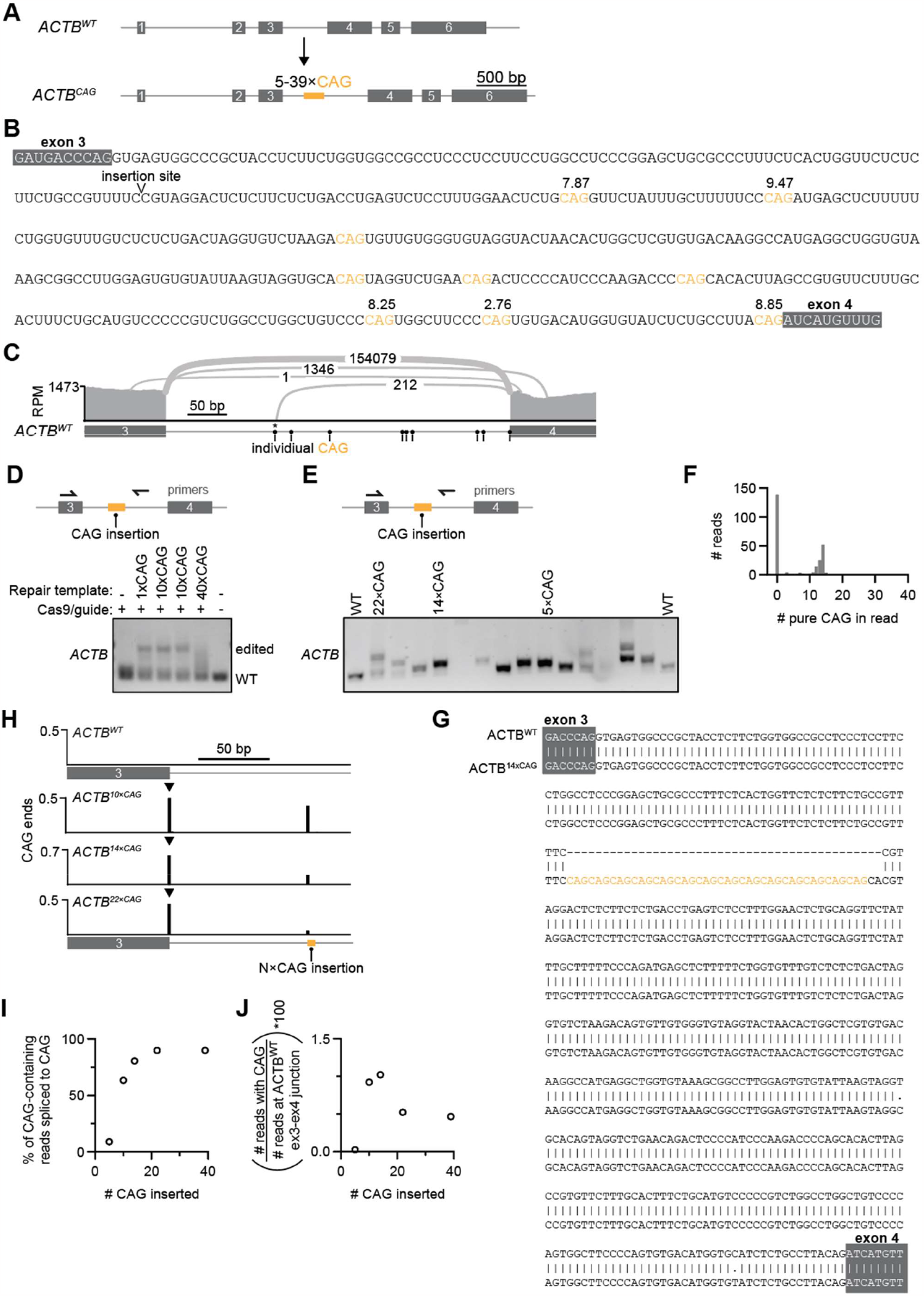
Insertion of a CAG repeat into an endogenous gene creates a new 3’ splice site. A. Schematic for CRISPR-mediated integration of various CAG repeat lengths into the intron 3 of *ACTB*. **B**. Complete sequence of *ACTB* intron 3. Individual CAGs are highlighted and 3’ MaxEnt scores are indicated for sites with positive scores. **C**. Sashimi plot for the *ACTB* intron 3 region. The y-axis represents the number of reads covering each base. Splice junctions are indicated as arcs between donor and acceptor, with the number of reads spanning the junction overlaid on the arc. The individual CAGs within the intron are indicated by circles. A single splice junction is marked with an asterisk: this site acts as a rare splice donor, not a splice acceptor. **D**. Endpoint PCR screen on polyclonal genomic DNA for knock-in of 120-base sequences with or without expanded CAG repeats to *ACTB* intron 3. Inserts had 1, 10, or 40×CAG repeats, and were padded to 120 bases with non-repetitive sequences (FLAG tags). An expanded 40×CAG repeat tract, but not short repeats, contracts when used as a ssDNA repair template. **E**. Representative endpoint PCR screen for individual clones of *ACTB*^*CAG*^ knock-ins. In total, >40 clones were screened to obtain a length series. Each clone has a different repeat length due to random truncations of the repair template during integration. **F**. Representative distribution for the number of CAG repeats in *ACTB* intron 3 across ∼200 Oxford Nanopore sequencing reads from genomic DNA of a single *ACTB* knock-in clone. About 60% of reads reflect the wildtype allele (0×CAG), while the remainder have ∼14×CAG. Some reads have fewer than 14×CAG, likely due to errors associated with nanopore sequencing. **G**. Sequence alignment for wildtype *ACTB* and *ACTB* with 14×CAG integration, which were generated from long-read sequencing of amplicons generated on the ACTB region. **H**. SATCfinder output for the *ACTB* intron 3 region in wildtype and CAG-inserted clonal populations. The wildtype *ACTB* region does not have any ≥3×CAG repeats. **I**. Quantification of the % of CAG repeat-containing reads that reflect splicing to the repeats. **J**. As panel I, but normalized to the number of reads spanning the wildtype exon 3 – exon 4 junction.

